# Artificial capillaries-on-a-chip with modular control over lumen size, architecture, in situ modifications and co-culture conditions

**DOI:** 10.64898/2026.01.29.702578

**Authors:** Arun Poudel, Joshua Edwin Nana Limjuico, Ujjwal Aryal, Md Tariqul Hossain, Saikat Basu, Pranav Soman

## Abstract

Currently in vitro models of microvascular biology rely on self-assembly of vascular cells in compatible gels. However, the stochastic nature of this process results in large variations in lumen sizes, perfusion continuity, and shear stresses making systematic and reproducible analysis challenging. Here, we report a new technology to generate artificial capillaries on a chip with custom control over lumen sizes and architectures using a combination of femtosecond laser cavitation and collagen casting within multi-chambered microfluidic chips. The design allows seeding of endothelial cells within capillary-sized microchannels and seeding of stromal cells within top-open silos, with independent control over seeding sequence and media compositions. Results show that endothelialized microchannels, coined as artificial capillaries, exhibit excellent barrier function with reproducible control over lumen sizes (ϕ=8-35µm) and their architectures (straight, curvatures, tapered, branched). The physical flow parameters measured across the lumen (namely, flow shear) and at the channel outlets (flow velocities) have been validated against high-fidelity numerical assessments from the Large Eddy Simulation scheme within the digitized versions of the microchannels. The experiment-computation compatibility enabled us to predict changes in regional velocity and wall shear stresses within artificial capillaries, for various capillary architectures. We also show that in situ editing of artificial capillaries in the form of adding new branches or adding occlusions is possible. Lastly, we developed a co-culture model which enables the study of stromal cells with artificial capillaries using conventional imaging methods. We envision that acellular chips with two seeding ports can be readily shipped worldwide and could potentially be adopted as a new technology to study microvascular biology in a reproducible manner.

## Introduction

Since capillaries are the primary site of nutrient exchange between tissue and blood circulation, understanding the mechanisms regulating capillary function in health and disease can be beneficial for development of new therapies. Inspired by angiogenesis and vasculogenesis processes, many in vitro models have been developed to study microvascular biology. Such models typically involve the use of endothelial cells within fibrin gels forming networks via self-assembly. However, the stochastic nature of capillary formation results in poor reproducibility with a wide range of lumen sizes (5-200μm), unpredictable perfusion connectivity, and locally varying fluid flow and shear stresses, which makes a systematic study about microvasculature challenging.^1–5^ To increase reproducibility, molding, bioprinting, and laser processing have been widely used to pre-defined endothelialized channels.^6–20^ The simplest method involves the removal of a tubular molds from polydimethysiloxane (PDMS) chips to leave behind a lumen (∼150–550μm) embedded within ECM matrix which is then lined with endothelial cells. To go beyond planar topology, 3D bioprinting has also been used, although each method enforces strict constraints on bioink composition. For instance, extrusion based direct printing requires high viscosity bioinks and rapid crosslinking mechanisms to print robust structures, while indirect printing requires the use of sacrificial materials that can be removed without generating any cytotoxic side effects, and light-based printing requires photo-sensitive and low-viscosity bioinks. At present, only multiphoton laser ablation or degradation can generate capillary-sized channels^21–25^, although negative impact on cell viability during laser scanning and endothelializing channels less than 50 μm lumen size remains difficult. To address these challenges, we report a new technology to generate highly reproducible artificial capillaries embedded within unmodified extracellular matrix (collagen). Here, we have user-defined control over lumen size (7-35µm), architectures (branching, tapers, zig-zag), in situ editing, and incorporating stromal cells during active cultures, yet enable easy-to-use cell seeding method, and be compatible with standard imaging and analysis methods.

## RESULTS

### Design and fabrication of chips to generate capillary-sized microchannels in 3D collagen

Briefly, a combination of 3D printed Master Molds, collagen casting, and femtosecond laser-assisted cavitation was used to generate capillary-sized microchannels embedded within collagen in PDMS chips. First, Projection Stereolithography (PSLA) was used to make customized three-chambered microfluidic devices using Polydimethysiloxane (PDMS) microfluidic devices. Here, digital light patterns (405 nm) modulated by Digital Micro-mirror Devices (DMD) was used to print a negative master mold using a photosensitive PEGDA (250kDa) resin. **(Fig. 1A, SI-1)** Replica molding was used to generate PDMS chips with three chambers seperated by an array of microposts chips **(Fig. 1B, SI-2).** The total height of the chip was 4 mm while the height for all three chambers in the chip was 200 µm. The final devices consist of three-chambers: a central chamber (Ch#2; 0.9 mm wide, designed to house crosslinked collagen) flanked on either side by two side chambers (Ch#1 and Ch#3; 1 mm wide, for media exchanges). In this work, we choose type I collagen as model ECM due to its abundance in *in vivo* tissues, and its wide use for developing 3D cell culture models. Before pipetting collagen solution within PDMS chips, chips were surface modified to prevent delamination of crosslinked collagen from the channel surfaces during active cultures. Then, type I collagen solution (5mg/ml) was gelled within the central chambers of UV sterilized chips. After 24 hours, a focused femtosecond laser (1.3 W, 40X water immersion objective, 0.8 N.A., 800 nm) was used to generate 3D microchannels architectures. To reliably generate 3D channels of defined lumen sizes, laser scanning was performed within collagen gel about 50µm above the bottom glass surface. **(Fig. 1C)** Reflectance microscopy images show top and cross-sectional views of microchannels embedded in collagen matrix. **(Fig. 1D, SI-3)** Laser dosage above 2 × 10^5^ Jcm^-2^ resulted in cavitation and formation of lumens of ∼3µm size. During laser scanning, the radial expansion of the bubble generates a shockwave which locally breaks down the collagen network and leaves behind a hollow lumen in its wake. Protocols were optimized to ensure repeatable and stable channel formation without collapse. At this speed, a single scan of 1mm takes ∼ 10 seconds. In a single scan, a lumen size of 3 µm can be generated while multiple scans with varying lateral offsets (1, 2 and 4µm) were optimized to generate lumen size from ∼3-40 µm. **(SI-3)** Based on the fabrication speed of 100 µm/s, a lumen of 20 µm required ∼16 scans, with a duration of 170 seconds while a lumen size of 35 µm required ∼28 scans, corresponding to around 298 seconds of fabrication time. The plot **(Fig. 1E)** shows the relationship between the number of laser scans, target lumen size, and total fabrication times. Lumen sizes, as measured by confocal reflectance microscopy across multiple chips, showed a high degree of reproducibility with close to circular cross-section. These laser-sculpted chips could be stored under hydrated conditions (PBS in side-chambers) for up to 3 months without any obvious effect of deterioration, although we used all chips within 2 weeks of fabrication.

**Figure 1.**
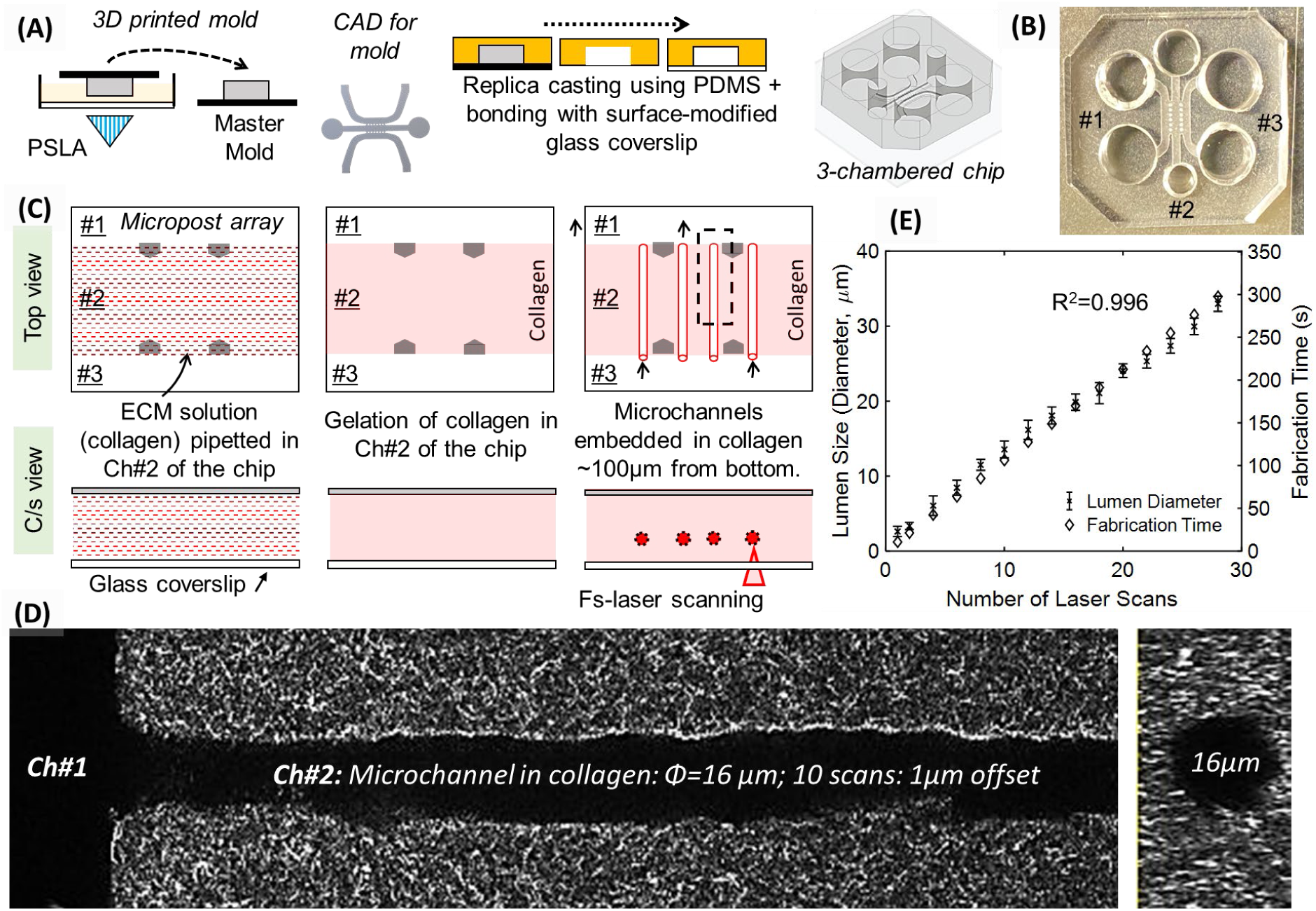
(A) Process flow showing the fabrication process to make PEGDA master molds and final PDMS device for chip design A. (B) CAD model of a three-chambered chip and a representative photograph of final PDMS chip bonded to glass coverslip. Photograph shows chamber 1-3 (Ch#1, Ch#2, Ch#3) separated by an array of microposts (C) Top and cross-sectional schematic views highlighting the process flow to make endothelialized microchannels within collagen (artificial capillaries) inside chips. (D) Confocal reflectance microscopy images of top and cross-sectional views of a representative microchannel embedded in collagen matrix. (E) Plot showing the relationships between target lumen size of microchannels, the number of laser scans, and total fabrication durations. Linear regression revealed a strong correlation (R²=0.996)

### Optimization of perfusion culture in chips (Figure 2)

Gravity-based flow was used to generate perfusion within microchannels embedded in collagen. To induce unidirectional flow, 1ml pipette tips were fitted to the inlet and outlet ports of chamber 1 and both pipettes were filled with equal volumes of PBS (or media during active cell-cultures) while the inlet and outlet ports of chamber 3 were left open. **(Fig. 2A)** Hydrostatic pressure gradient across the collagen gel, corresponding to the PBS height in pipettes, initiated a range of velocities in microchannels. To calculate the velocities within the microchannels, chips with 6 microchannels of lumen size 40 µm were developed. Different volumes (0.05 – 1ml) of microbead solution (yellow-6.0µm) was filled in the pipette tips, and time-lapse movies of bead movement was recorded (Hamamatsu, 40 fps). A region of interest, inside the lumen in chamber 2 was identified (**Fig. 2B**, dashed yellow box), and velocity measurements were averaged across five individual beads in each of the 4 different channels, across three independent samples. Bar plot shows that bead velocity decreased with decrease in pressure head. **(Fig. 2C)** Table shows that by simply changing the media volume, we can achieve a shear stress range between ∼10 to 65 dynes/cm² **(Fig. 2D)** close to the in vivo range reported in the literature (5 to 95 dynes/cm²)

**Figure 2.**
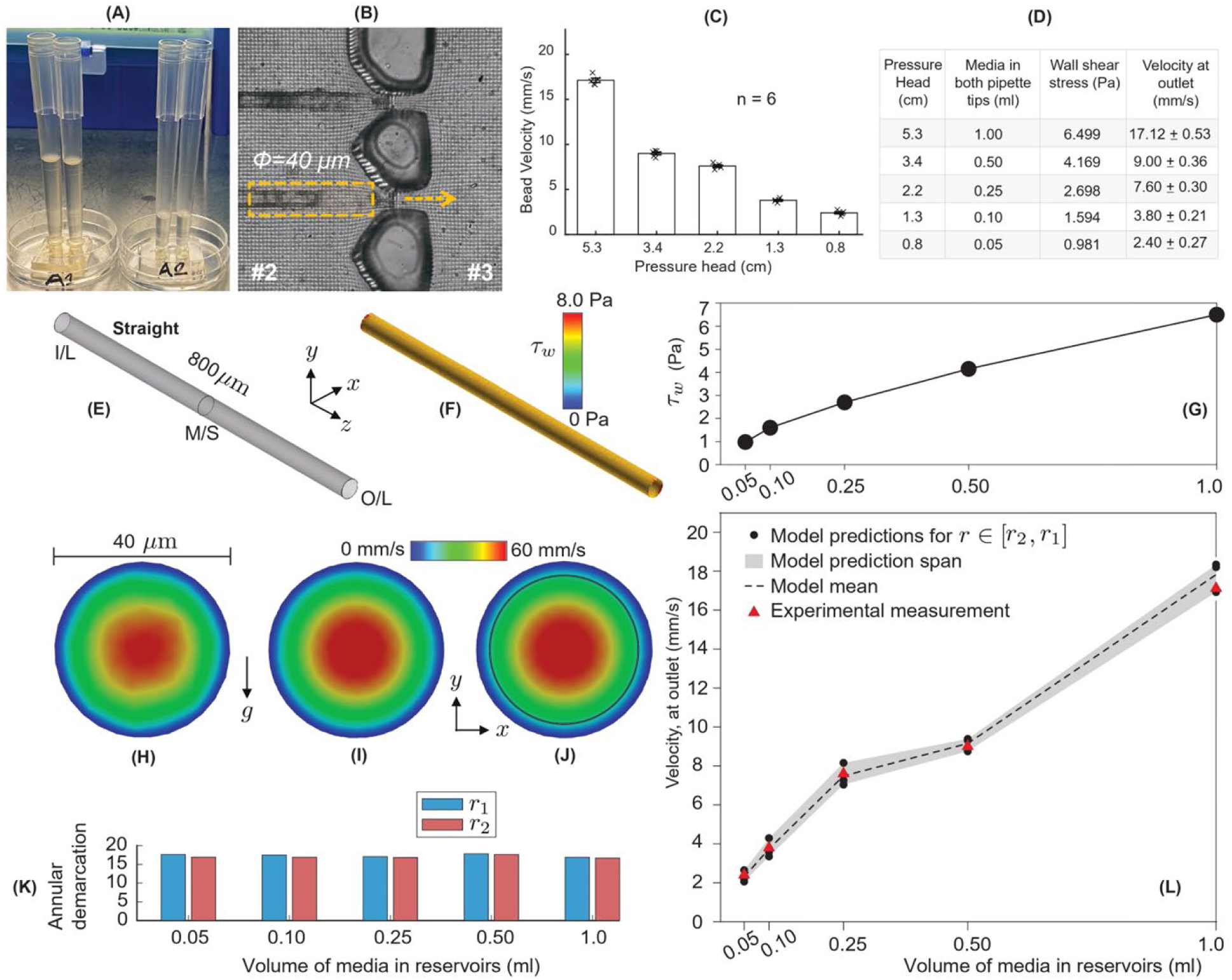
(A) Photograph of hydrostatic pressure driven perfusion setup showing 1ml pipette tips inserted in the inlet and outlet ports of Ch#1 while other ports are open to atmospheric pressures; this enables directional flow of media from Ch#1 to Ch#3 through microchannels embedded within collagen in Ch#2. (B) Brightfield image showing two microchannels (lumen size = 40µm) at the interface of Ch#2 and Ch#3 while dashed box highlights the Region of Interest (ROI) used to calculate the velocity of microbeads. (C) Plot and table shows the relationships between pressure-head, bead-velocities, and calculated wall shear stresses generated within the microchannels. (E). Digitized straight microchannel, showing the identified locations for inlet (I/L), mid-point section (M/S), and the outlet (O/L). (F). Surface map of the area-weighted average wall shear with the simulated pressure head at 5.3 cm. (G). Simulated wall shear values, for the tested range of pressure heads (labeled in terms of the volume of media in the pressure head reservoirs). (H-J). Simulated velocity fields, respectively at the inlet, mid-point, and outlet of the digitized microchannel (per panel E). (K). Data on the annular region location in the outlet field, bounded by r1 and r2, commensurate with the experimentally measured bead velocities. (L). Experiment-computation comparison for velocity measurements at the microchannel outlet for the tested pressure heads.

Due to scattering induced by the collagen matrix, it was challenging to measure bead velocities for smaller lumen sizes, so we developed new computational models to calculate the velocities and wall shear stresses. We also numerically simulated the pressure gradient-driven flow physics within a digitized replica of an isolated straight microchannel (**Fig. 2E**), spatially segregated into approximately 1.7 million graded, unstructured tetrahedral elements. The mesh resolution within the in silico domain was carefully derived from a grid sensitivity analysis of the simulated flow outcomes (see **SI-4**). For reliable computational fluid dynamics (CFD) estimation of the transport parameters, we implemented the high-fidelity Large Eddy Simulation (LES) scheme, with the dynamic kinetic energy transport model resolving the subgrid-scale fluctuations (the simulation time-steps were 0.0001 s; see ^26^ for related mathematical formalism applied in a different micro-scale physiological system). The ambient pressure conditions in the simulations were guided by the corresponding experimental data (so, separate simulations were developed with pressure heads of 5.3, 3.4, 2.2, 1.3, and 0.8 cm – applied from inlet I/L to outlet O/L; see **Fig. 2E**). The simulated area-weighted average wall shear for the 5.3-cm pressure head was 6.45 Pa; the surface map is shown representatively in **Fig. 2F**. We have subsequently plotted all the simulated wall shear averages for the different pressure head conditions (see **Fig. 2G**); the specific values are *τ_w_* **=** 6.45, 4.15, 2.69, 1.59, and 0.98 Pa – respectively for the highest-to-lowest pressure heads. The trend matches with the experimental assessments for shear stresses (based on the measured velocity gradients); see table in panel **2D**. The simulated velocity fields are next shown in **Fig. 2H-J**; therein note the narrow black annular region in the outlet field in **Fig. 2J**. The experimental data (see table) indicated bead velocities of *v_e_* = 17.12, 9.0, 7.6, 3.8, and 2.4 mm/s at the microchannel outlet (respectively under pressure heads of 5.3, 3.4, 2.2, 1.3, and 0.8 cm).

The corresponding standard deviations on the velocities were *σ* = 0.53, 0.36, 0.30, 0.21, and 0.27 mm/s. From Hagen-Poiseuille equation, we estimated the annular region extents in the simulated outlet field (e.g., in **Fig. 2J**) where the velocities stay within *v_e_* ± *σ*; for instance, for the 5.3-cm pressure head, the annulus is bound by *r_1_* = 17.65 mm and *r_2_* = 16.59 mm (the comprehensive set of {*r_1,_ r_2_*} all pressure gradients is shown in **Fig. 2K**). The finding conforms with the expectation for the bead locations to be skewed toward the channel walls during the experimental measurements (owing to the diverging nature of the flow streamlines at the exit cross-section). From the numerical grid points within this annulus, we extracted the simulated velocity values; for the 1.7-million element mesh, there were three such grid points within the narrow annulus. The validative comparison between the simulation-derived channel outlet velocities and the experimental measurements (with red markers) is demonstrated in **Fig. 2L**.

### Endothelialization of capillary-sized channels embedded within collagen matrix

Model vascular endothelial cells (HUVECs), cultured under standard conditions, were seeded in chambers 1 and 3 at a concentration of 0.5M cells/mL and 24 hours later, perfusion flow was initiated using 1ml media in pipettes tips. Brightfield images show that cells readily migrate within the channels within 24-48 hours and by Day 4 microchannel surfaces are completely covered by HUVECs. Sometime, when a seeding concentration of 1M/ml or higher is used, channels can be blocked (arrow, **SI-5**), which can be removed by simply reversing the media flow by adding media-filled pipetted to the other side-channel ports. For seeding concentrations of 0.5M/ml or less, no blockages were noted, and unidirectional media flow was performed.

Morphology of cells, characterized by staining for f-actin, show uniform coverage of cells within channels of lumen 35µm. **(Fig. 3A)** Cross-sectional images show that lumens are open throughout the length of the channels. To characterize the patency and barrier function of these channels, FITC-dextran (70kDa) solution was used on chips with microchannels lined with or without HUVECs. **(Fig. 3B, SI-6)** Results show that HUVECs prevent diffusion of dye into the surrounding collagen matrix. In fact, since the permeability for HUVEC-lined channels was undetectable with 70kDa dye, we repeated this experiment with a lower-molecule weight FITC-Dextran 4kDa dye. Even the permeability coefficient for 4kDa (0.3 × 10⁻^5^ cm/s, **V:1**) was found to significantly lower to 2.54 × 10⁻^5^ cm/s obtained for control collagen channels (with 70kDa dye, **V:2**). This shows that endothelialized channels have robust barrier function.

**Figure 3.**
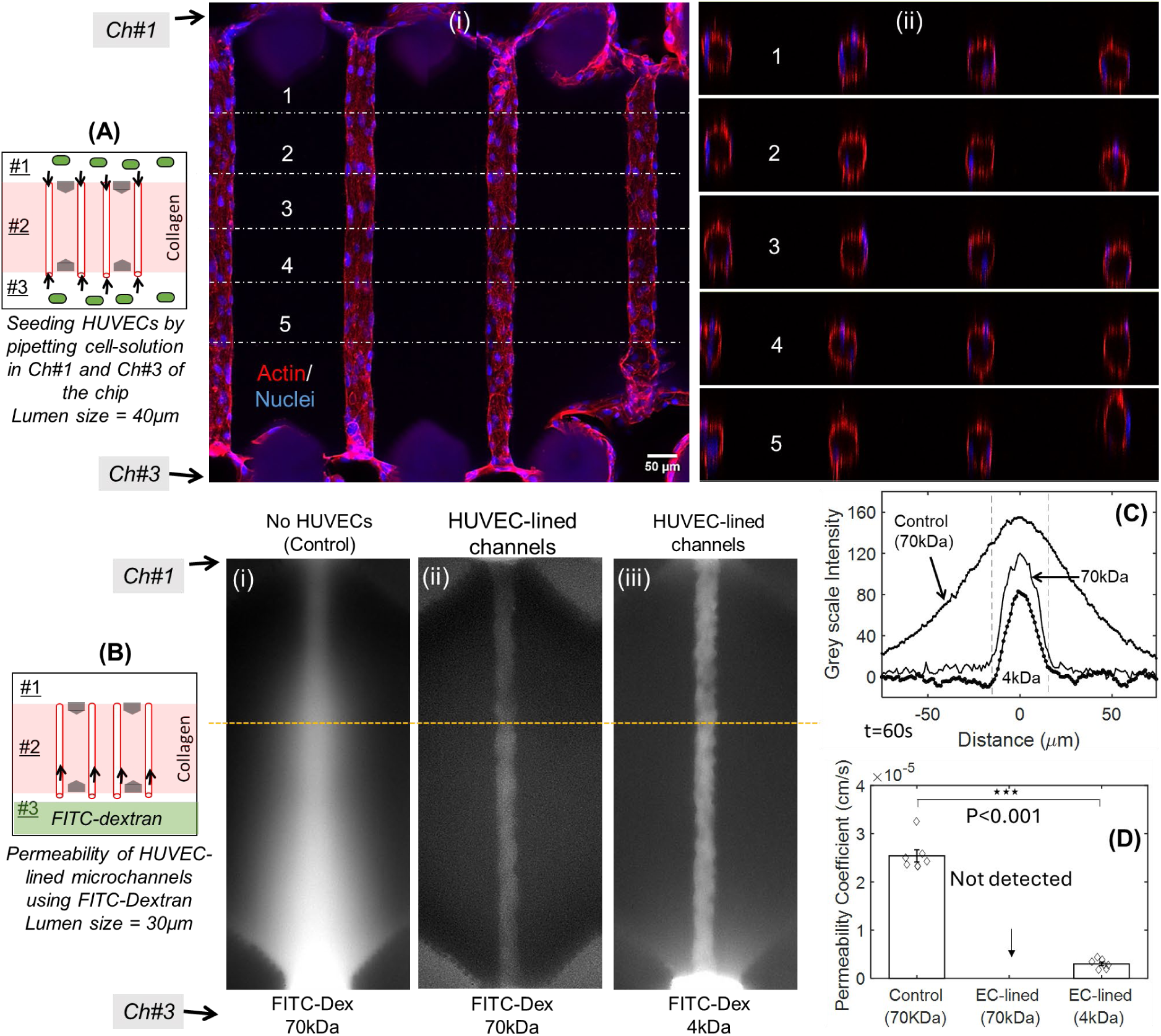
(A) (i) Schematic showing seeding of HUVECs in Ch#1 and Ch#3 of the chip to generate endothelialized microchannels (diameter=40µm) embedded in collagen ∼50µm above the bottom glass coverslip. (i-ii) Representative fluorescence microscopy top and cross-sectional views highlighting cell morphology on Day 5. (B) Schematic showing the process flow for testing barrier function of artificial capillaries. (i-iii) Representative fluorescence microscopy images showing diffusion of FITC-dextran from the microchannels into the collagen under various conditions. (C) Fluorescence intensity plots comparing leakage of dye from the microchannels, (D) Permeability coefficient of artificial capillaries.

### Characterization of endothelialized lumens

Chips with HUVEC-lined microchannels were stained with CD-31, a marker for Platelet/Endothelial Cell Adhesion Molecule-1 (PECAM-1), which is known to play a role in regulating vascular permeability. Fluorescence images show that artificial capillaries (ϕ∼25µm) remain open and exhibit monolayer of cells around the internal surfaces of the channels. **(Fig. 4A, B, V:3,4)** From the cross-section of endothelialized microchannels, we found 1 to 3 nuclei were used to form the lumen based on the location within the microchannels. **(V:5-7)** These structures resembled the cross-section of *in vivo* capillaries, with thin, elongated nuclei with slight bulges projecting into the lumen. **(Fig. 4B, iii, SI-7)** We analyzed the changes in cell areas, thickness of the cytoplasm, and orientation of nuclei with reference to the direction of media flow. **(SI-8)** with Based on VE-cadherin staining, artificial capillaries exhibited an elongated cobblestone-like morphology, typically seen under unidirectional flow conditions. **(Fig. 4C)** For Golgi apparatus, no polarization along the direction of flow was observed. Assuming complete coverage of HUVECs in the microchannels, based on our permeability results, we divided the total area of the cylindrical channel by the number of nuclei to get a metrics of cell spread in channels with various lumen sizes. We found that, as lumen size decreases, cells spread more; a smaller number of cells can spread and wrap with large curvature smaller lumen channels (10 µm vs 40 µm). **(Fig. 4D).** The thickness of the cytoplasm for each lumen size was measured. Results show that for lumen size of 10 µm, a minimum thickness of 0.6 µm was recorded, while a maximum of 1.8 µm was calculated for larger lumen sizes (20-40 µm). **(Fig. 4E)** Other than a lumen size of 25µm, all other lumen sizes show a nuclear orientation perpendicular to the direction of flow. **(Fig. 4F)**.

**Figure 4.**
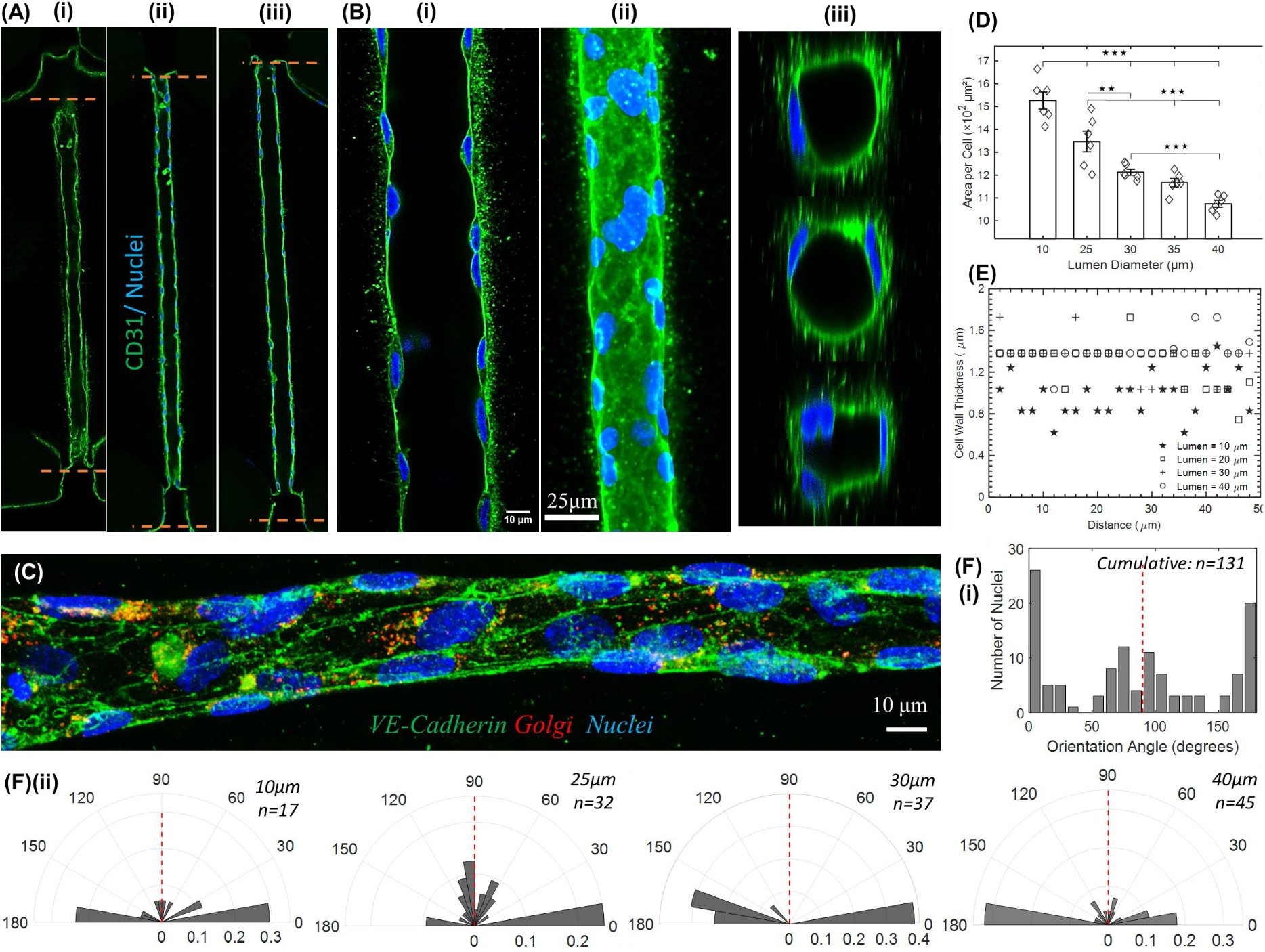
(A) Representative images showing (expression of CD-31 (PECAM-1, green). ϕ∼25µm. (B) High resolution images at a single-plane (i), z-stack composite (ii), and cross-sectional images along the length of the microchannel (iii) (C) Images showing staining of VE-cadherin (green), Golgi apparatus (red), and nuclei (blue). Plots showing variations in cell area (D) and cell thickness (E) as a function of lumen size. (F) Plots showing changes in nuclei orientation with respect to flow direction.

### Customization of microchannel architecture

To mimic the complexities of *in vivo* capillaries, we generated user-defined customized channel topologies followed by HUVEC seeding, as described earlier. Brightfield images show the migration of HUVECs in microchannels of different geometries (Day 0 – 4, **SI-9**) Custom designs included from the top **(Fig. 5A i, ii)**: (i) branched, (ii) zig-zag, (iii) tapered, and (iv) straight. Brightfield images show the formation of HUVEC-lined channels. On Day 5, permeability was characterized. Cross-section of branched topology shows open lumen with nuclear morphologies similar to straight channel topology. **(Fig. 5B)** Permeability studies, performed on HUVEC-lined channels using FITC-Dextran (4kDa) on Day 5, show that topology of the channels does play a role in their permeability properties. **(Fig.5Ci)** Some local regions of leakages can be seen in the branched and possibly the zig-zag lumen geometry as opposed to straight lumens that show no leaks. Permeability characterization show that branched (4.5 × 10⁻⁶ cm/s), zig-zag (3.237 × 10⁻⁶ cm/s), and tapered (3.410 × 10⁻⁶ cm/s) topologies show higher permeability as compared to straight lumen geometry (2.441 × 10⁻⁶ cm/s).**(Fig.5Cii)** Additionally, the computationally simulated velocity fields (exemplified by 30 representative fluid streamlines) and the resulting area-weighted average of the wall shear – for the two channel shapes highlighted in **Fig.5Bii** – are included in **Fig. 5D-G** (for the zig-zag channel shape) and **Fig. 5H-K** (for the branched channel shape). The cross-sectional diameters of the in silico domains were maintained consistently at 35 µm (in conformity with the physical design). Within the zig-zag channel, the four panels in **Fig. 5E** demonstrate the velocity fields at the cross-sections marked by dashed lines in **Fig. 5D**, in the same left-to-right order. The two panels in **Fig. 5G** correspondingly show the strain rate contours within the bulk for the cross-sections marked by dashed lines in the wall shear demonstration in **Fig. 5F**, in the same left-to right order. Similarly, within the branched channel, the three panels in **Fig. 5I** demonstrate the velocity fields at the cross-sections marked by dashed lines in **Fig. 5H**, in the same left-to-right order. The panels in **Fig. 5K** show the strain rate contours in bulk for the cross-sections marked by dashed lines in **Fig.5J**’s wall shear map, in the same left-to right order. As expected, the shear effects maximize at the corners of the channel geometries. The simulations are, however, representative, being only for the pressure head 5.3 cm (in this context, note the broader range of pressure heads tested in **Fig. 2** in the simpler channel).

**Figure 5.**
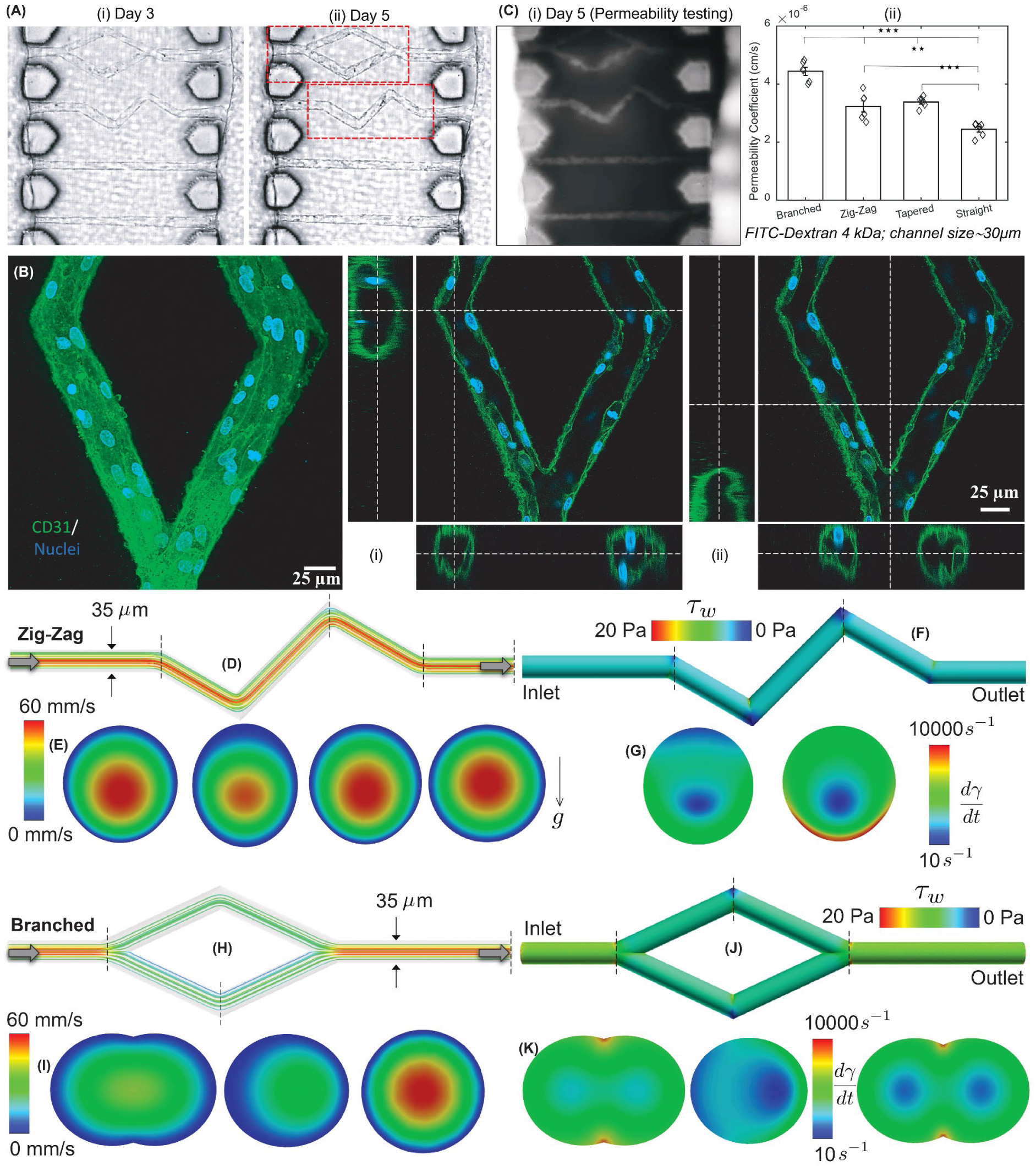
(A) Representative brightfield images showing artificial capillaries with custom geometries (branched, zig-zag, straight, tapered). (B) Cell morphology via PECAM staining within branched geometry capillaries. Lumen size=30um (C) Fluorescence image during permeability testing (i) and its quantification (ii). While arrows point to regional leaks in areas of high shear stresses. D. Numerically simulated velocity streamlines in the zig-zag channel shape highlighted in panel 5Bii. (E). Cut-aways in the velocity field for the cross-sections marked by the dashed lines in panel D, in the same left-to-right order. (F). Numerically simulated wall shear in the zig-zag channel. (G). Strain rate maps for the cross-sections marked by the dashed lines in panel F, in the same left-to-right order. (H). Numerically simulated velocity streamlines in the branched channel shape highlighted in panel 5Bii. (I). Cut-aways in the velocity field for the cross-sections marked by the dashed lines in panel H, in the same left-to-right order. (J). Numerically simulated wall shear in the branched channel. (K). Strain rate maps for the cross-sections marked by the dashed lines in panel J, in the same left-to-right order.

For the zig-zag channel, the simulation-derived mean outlet velocity and wall shear stress were 22.86 mm/s and 5.13 Pa, respectively. The overall range of wall shear (*τ_w_*) was 0 – 20.37 Pa, with the strain rates (dγ/dt) in the bulk ranging between 17.01 – 16596.95 s^-1^. The latter could be used to compute the range of bulk shear *τ* = µ(dγ/dt), with the numerically implemented viscosity coefficient being µ = 0.001003 Pa.s (for water). Accordingly, in bulk, *τ* ∈ (0.017, 16.646) Pa. Next, for the branched channel, the corresponding estimates for the simulation-derived mean outlet velocity and wall shear stress were respectively 31.63 mm/s and 4.98 Pa. Therein, the overall range of wall shearwas 0 – 11.41 Pa, with the strain rates in the bulk ranging between 15.56 – 11113.39 s^-1^. Following the same computation as above, the shear force in bulk turns out to be *τ* ∈ (0.016, 11.147) Pa.

### In situ editing of artificial capillaries by adding new branches

Due to external stressors, injury or disease, capillary network reconfigures itself and alters the distribution of blood flow. To test whether such structural rearrangement can be induced within engineered capillaries, laser was used to remove collagen from target section to generate new microchannel connection between adjacent capillaries. Here, HUVEC-lined channels were generated in the chip, and on Day 5, a femtosecond laser was used to create connections between two HUVEC-lines microchannels. **(Fig.6Ai,ii, SI-10)** Red arrows show the presence of new microchannels that connect two adjacent HUVEC-lined microchannels, while red circle shows the generation of an incomplete channel, which is only open on one side. Brightfield snapshots show zoomed-in images of a liquid droplet flowing through the newly generated connection. **(Fig. 6B)** Within 24 hours, cells migrate from the existing HUVEC-lined channels into the newly formed microchannel and generate a new branch in the network in 2-3 days. Confocal images show expression of PECAM while f-actin and nuclei morphology confirm the presence of open lumen and cell coverage within new branches. **(Fig. 6C)** In one of the branches, marked by a circle, incomplete ablation of channel resulted in the creation of a dead-end. In this case, HUVECs migrated inside the channels but retracted soon after resulting in a failed connection. In another experiment, GelMA (5%) was perfused into an engineered capillary, and an occlusion was generated by two-photon crosslinking; this decreases the lumen size from 20µm to ∼5 µm. **(SI-11)** Perfusion of FITC-dextran 2000kDa confirmed the presence of this partial occlusion, where flow can be seen to go around the partial occlusion.

**Figure 6.**
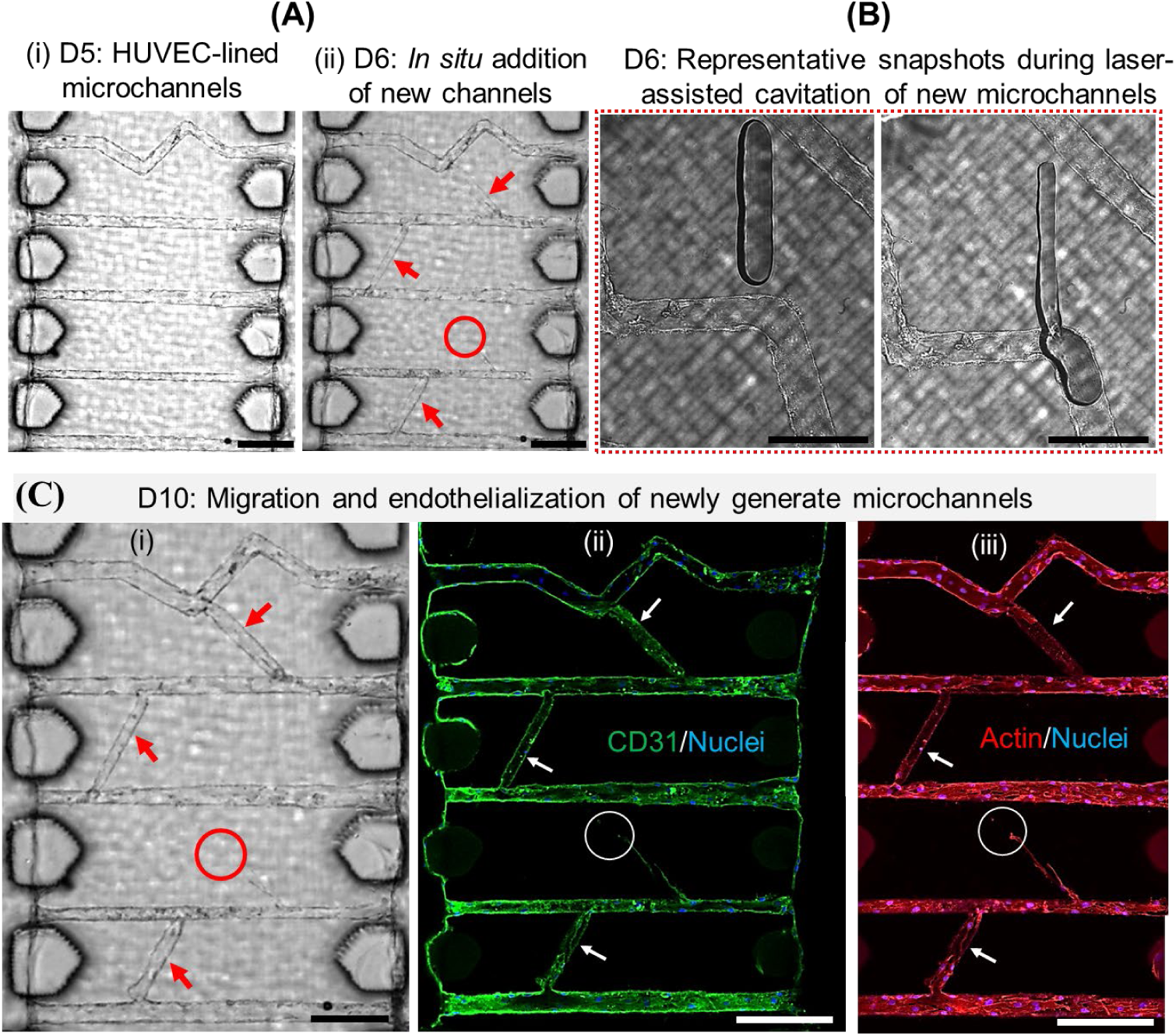
(A) Brightfield images showing addition of new branched to previously endothelialized microchannels on Day 6 (D6). (B) Zoomed-in sequence of 2 images showing media flows through newly generated branch. (C) Brightfield (i) and fluorescence microscopy (ii-iii) images showing endothelialization of new branches after 4 days. Scale bar A: 200 μm; B: 50 μm; C: 200 μm

### Co-culture of model stromal cells with artificial capillaries embedded in collagen

To enable co-culture of endothelialized channels with model stromal cells (fibroblast-like 10T1/2s), we designed a new chip (design B). In this design, top and bottom parts, printed via projection stereolithography (PSLA), was assembled and replica casted using PDMS to fabricate three-chambered chips with a pair of silos in the central chamber (Ch#2). **(Fig. 7A, SI-12-15, V:8)** A conical plug with two pins (diameter=150 µm), printed using PSLA, was fitted into the central chamber of the PDMS chip, to ensure a vertical gap of ∼50 µm between the base of the pins and the bottom glass coverslip. **(Fig. 7B, SI-16, 17, V:9)** Then, rat-tail type I collagen solution (5mg/ml) was perfused in chamber #2 and allowed to gel at 37°C for 30 minutes. Post gelation of collagen, plug was removed to leave behind two cylindrical silos (∼150 µm) embedded within gelled collagen in chamber 2. **(Fig. 7C)** The plug-pin design is such that a larger reservoir (∼1mm, shown in **Fig.7Avii**) was generated to enable easy pipetting of media and cell solutions. Femtosecond laser scanning was used to generate a pair of microchannels (∼30 µm lumen size) on either side of central silos. Reflectance confocal microscopy image shows both the central silo (∼150 µm) alongside one of the microchannels at ∼50-100 µm inside gelled collagen matrix in chamber 2 of the chip. **(Fig. 7D)**

**Figure 7.**
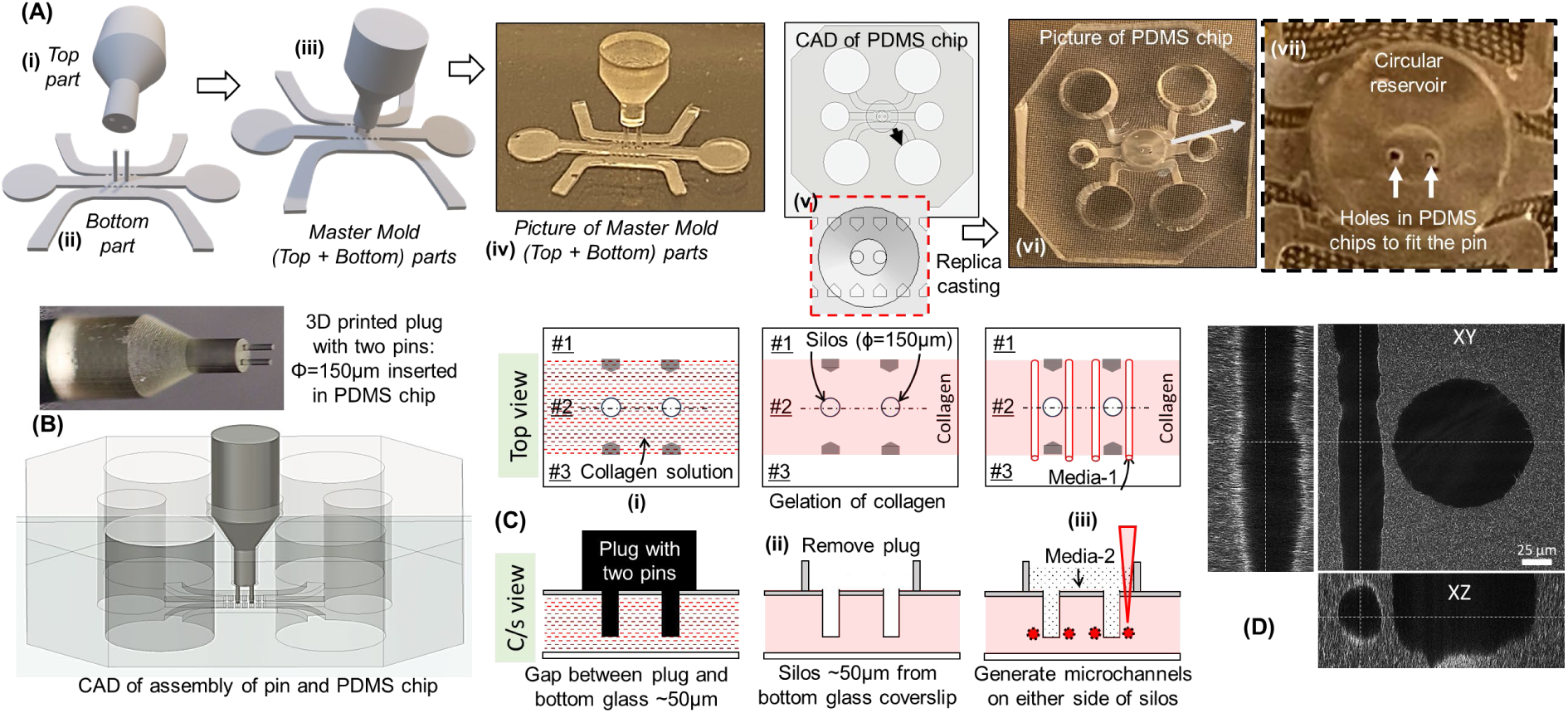
(A) Chip design B to facilitate co-culture experiments via simple cell-seeding. (A) Schematic showing the process flow: Master mold is generated by assembling the top and bottom parts, then replica casting is used generate a three-chambered PDMS chip (iv-v) with a circular reservoir with two holes (vi-vii). (B) 3D printed plug (photography) with two pins inserted in PDMS chip (CAD assembly). (C) Schematic showing the process flow starting with perfusion of collagen solution in Ch#2, and post-gelation, pin is removed leaving behind silos. Then fs-laser cavitation is used to generate microchannels on either side of the silos. (D) Confocal reflectance microcopy images show the microchannel and silo.

HUVEC solution (0.5M/cells) was pipetted in the side chambers of the chip to enable endothelialization in 2 days. On Day 3, model stromal cells (fibroblasts like 10T1/2s, 0.5M/cells) were pipetting into the open reservoir on top of chamber 2. Brightfield images show migration of fibroblasts into the collagen matrix within 24 hours and surround the endothelialized microchannels within 6 days. **(Fig. 8A, SI-18)** Tiled confocal images showing fibroblasts migrating from central silo (CD44, yellow) towards HUVEC-lined microchannels (CD31, green). By Day 6, fibroblasts are seen wrapping around the lumen from the outside. **(Fig. 8B, SI-19, V:10,11)** For some chips, we observe the fibroblasts migrate near the HUVEC-lined microchannels and actively decrease the lumen sizes **(Fig. 8C I,ii)**, while in some other chips, we no change in lumen sizes was observed. **(Fig.8Ciii)** Next, we repeated the same experiments with lumen size of ∼8-12 µm. Here, microchannels were generated such as their lumen size change cross-section from lumen size of 20 µm to 8µm in the vicinity of the central silo. **(Fig. 9A)** While the lumen remains open, confocal images show that lumen size varies between 8-12 µm as fibroblasts interact with HUVEC-lined channels. **(Fig. 9B,C, V:12-15)** VE-cadherin staining shows an elongated morphology possibly due to the ablated template as well as unidirectional flow conditions. **(Fig. 9D)**. We have also numerically simulated the transport trend in an in silico idealization of this channel shape. In this configuration, the inlet and outlet diameters were maintained at 28 µm, while the diameter progressively decreased to 8 µm at the mid-section. Under a simulated pressure head of 5.3 cm, the mean velocity at the outlet and the wall shear stress were estimated to be 0.95 mm/s and 0.84 Pa, respectively (see **Fig. 9E**). The cross-sectional views of the velocity fields are included in **Fig. 9F** (at the tapered center) and **Fig. 9G** (at the flow outlet). **Fig. 9H** demonstrates the wall shear trend (maximizing at the tapered region), with strain rate fields in bulk shown in **Fig. 9I** (at the center). Herein, the maximum and minimum wall shear stress values are 17.857 and 0 Pa, respectively. The maximum and minimum strain rate values are 13866.280 and 9.668 s^-1^. Panels **F**, **G**, and **I** are on the same scale.

**Figure 8.**
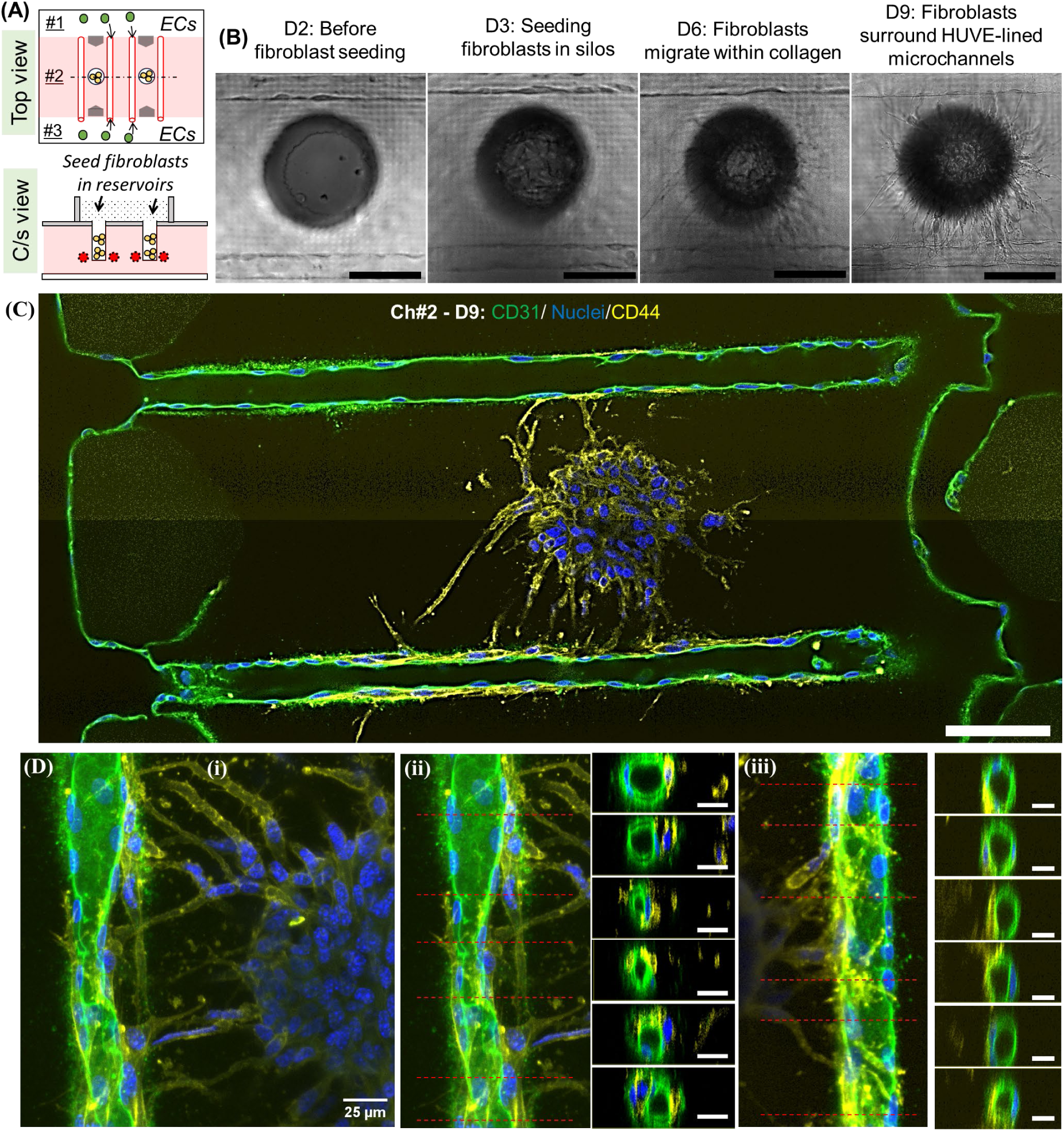
(A) Schematic showing co-culture of model stromal cells (fibroblasts-like 10T1/2s, gold color) in silos in close vicinity of HUVEC-lined microchannels (lumen size = 30µm). (B) Representative brightfield images showing addition of fibroblasts in silo on Day 3, followed by their migration into collagen matrix, and wrapping around artificial capillaries. (C) Titled confocal images showing the entire chip on Day 9. Image taken from the z-plane approximately at the center of the microchannels. (D) Confocal images of top and cross-sectional views from another chip showing stromal cell interactions with artificial capillaries (green). Scale bar B-C: 100 μm; D: 25 μm

**Figure 9.**
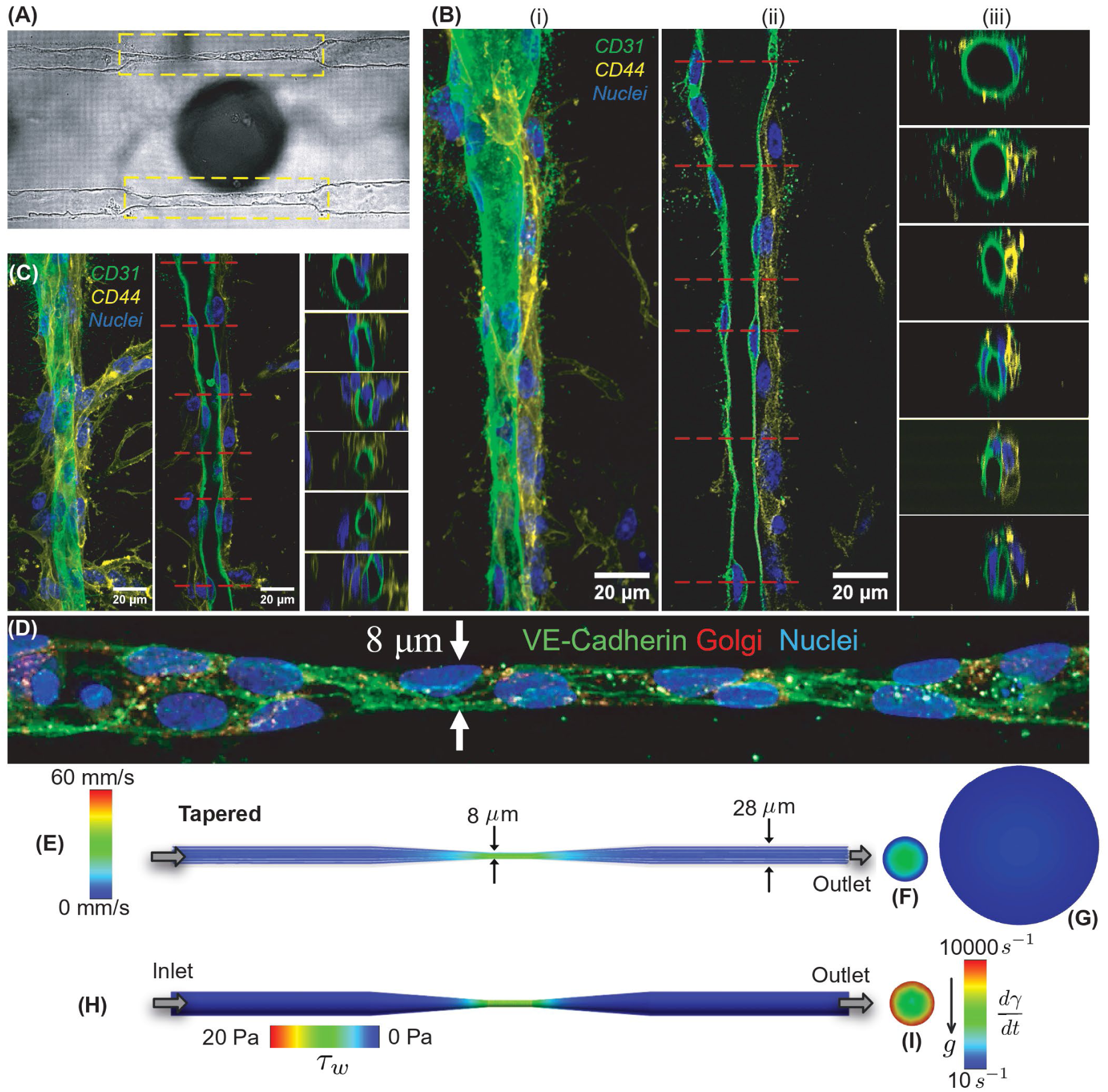
(A) Brightfield images of HUVEC-lined microchannels with change in lumen cross-section from 28µm to 8µm closely associated with a central silo. (B-C) Representative z-stacked, single-plane, and cross-sectional images showing fibroblasts (yellow) interacting with endothelialized microchannels (green). (D) Representative images showing staining of VE-cadherin (green), Golgi apparatus (red), and nuclei (blue). (E). Numerically simulated velocity streamlines for an idealized mid-tapered channel, mimicking the shape from panel D. The inlet and outlet diameters in the computational domain were maintained at 28 µm, while the diameter progressively decreased to 8 µm at mid-section, (F-G). Cut-aways in the velocity field at the middle point of the channel and at the outlet, respectively. (H). Numerically simulated wall shear in the tapered channel. (I). Strain rate map at mid-section. Panels F, G, and I follow the same length scale.

## Discussion

Capillaries, critical for nutrient exchanges between blood and surrounding tissues, can vary widely based on their location within the body. For instance, continuous capillaries (lumen size ∼5–10µm) are found in organs where selective exchanges are crucial (blood-brain barrier) while sinusoidal capillaries (lumen size ∼30-40µm), are found in organs that require the passage of larger molecules (liver, bone marrow). Moreover, drastic changes in capillary properties are often found in the same tissue. For instance, distinct capillary types were identified in long bones of mice ^27–29^ where oxygenated arterial blood is preferentially fed into type H columnar (straight) capillaries (∼10-15μm in diameter) followed by type L highly interconnected sinusoidal (branched) capillaries (∼20-30μm in diameter), before flowing into larger veins. Like capillary size and topology, their lengths could also vary widely between 50-500 µm. Since capillary-stromal interactions play a vital role in tissue development, aging, and diseases, having the ability to reproducibly generate capillaries on a chip is vital for the discovery of unknown crosstalk mechanisms between capillaries and organotypic function.

At present, the ‘gold standard’ to make capillary on a chip rely on self-assembly of vascular endothelial cells within relevant matrix (fibrin, collagen)^30–32^, which results in randomly organized microvasculature with wide variations in lumen sizes, architectures, perfusability, and associated shear stresses. To address these issues, a wide range of in vitro models have been developed to study microvasculature biology. Most common models cast ECM gels around sacrificial molds (needles, stamp, patterns) to generate microvessels of defined sizes.^3, 33, 34^. Despite its wide use in the field, capillary-sized lumens and complex topologies remain challenging with this method. Advances in extrusion bioprinting has made it possible to generate 3D channel topologies using a wide range of printable sacrificial materials.^35–37^ Other methods like electrohydrodynamic inkjet bioprinting can achieve a lumen size of 30µm^38^, although low-viscosity bioinks result in less stable structures. Manual removal of melt electro-written PCL fibers from cell-laden hydrogels can generate lumen sizes from 10-41μm,^39^ although branched or curved topologies cannot be realized. Light-based methods such as laser tweezers, stereolithography, digital light processing, and volumetric bioprinting, have also been developed to generate complex microvascular structures,^40–44^ although only multi-photon laser processing is capable of generating capillary-sized channels.^45–50^ Moreover, challenges of media perfusion and endothelialization within capillary-sized lumens and incorporation of supporting cells in relevant ECM around the lumens remain to be solved. In this context, we report a new technology to generate artificial capillaries on a chip with custom control over lumen sizes and architectures using a combination of femtosecond laser cavitation and collagen casting within multi-chambered microfluidic chips.

We show that femtosecond laser assisted cavitation can generate microchannels embedded within natural collagen matrix with defined lumen size range (8-35µm). We do not see any contraction of collagen channels after seeding HUVECs, as reported by other groups^6, 50^, possibly due to small cross-section of the microchannels and higher collagen concentration (5mg/ml) used in this work. We found that collagen below 2.5mg/ml results in unstable microchannels implying a stability threshold is necessary to generate structurally robust channels. Since mechanisms that govern laser-collagen interactions to generate a specific lumen size are still not completely known, we relied on trial-and-error methods to optimize the lumen sizes in this work. Although we focused on making in-plane microchannels ∼50-100µm inside collagen matrix, 3D out-of-plane channels are also possible by programming the stage accordingly, as shown in our previous work.^51^ Processing of collagen within PDMS chips made using 3D printed molds offer fast processing times and design flexibility. In the future, the processing time can be substantially decreased by using a galvanometric scanner instead of a scanning stage used in this work. We chose easy to culture, inexpensive, and widely used HUVECs, as a model cell to allow direct comparisons with existing model systems, although our method can be agnostic of both ECM and cell types. To demonstrate this capability, we synthesized hyaluronic acid methacrylate (HAMA) hydrogel and photo-crosslinked it within the central chamber of the chip followed by laser ablation of microchannels of varying lumen sizes (3-40µm).**(SI-20)** Unlike protein-based collagen, hyaluronic acid is sugar-based component found in ECM. Next, we replaced HUVECs with Endothelial Colony Forming Cells (ECFCs) and show that engineered capillaries can be generated with no change in our protocol. ECFC are specific type of endothelial progenitor cell with a high proliferative capacity to form new blood vessels *in vivo*, and more importantly they are clinically accessible source of autologous ECs.^52^ **(SI-21)**.

In this work, direct seeding of HUVECs in side-chambers for 24-hours followed by gravity-based perfusion culture for 48 hours resulted in robust capillary-sized vessels in 3 days. This is in contrast to reports that require a larger parent vessel before small-sized vessels can be grown.^46^ We found that HUVECs preferentially align within the lumens rather than migrating into the surrounding matrix, similar to what others have observed in other hydrogel systems^53^. Since our longest duration was only 12 days, more work is needed to test the influence of lumen size and curvature, fluid flow type (static, unidirectional, bi-directional) and the functional expressions of key biomarkers for longer culture durations. In our work, for lumen sizes (∼30µm), the lengths were the width of the chamber 2 (∼800µm) – an aspect ratio of ∼23, while for the smallest lumen sizes (∼8µm), the aspect ratio was ∼20. Further work is needed to study the effect of aspect ratio of artificial vessels and their long-term stability. The barrier integrity was initially tested with 70 kDa FITC-Dextran (close to albumin in terms of molecular weight, roughly a physical size of ∼100nm). Since the permeability coefficient for HUVEC-lined lumen was undetectable, we switched to a smaller size dye (4kDa FITC-dextran; similar to the size of small protein or peptide, ∼ 1.5nm), and even this showed a significant drop in permeability as compared to the controls that were performed with 70kDa dye. Our values are in range reported in the literature for *in vitro* HUVEC micro-vasculature models and close to *in vivo* microvessels.^54, 55^ We found that when the lumen size decreased, less number of HUVECs spread within the lumen; this is similar to other reports,^56^ which used large-scale volume electron microscopy data from mouse cortex to show that microvessels in transitional zones near penetrating arterioles and ascending venules are larger and typically composed of 2 to 6 endothelial cells, while capillaries intervening these regions are constructed from either 1 or 2 endothelial cells. We characterized the endothelial vessel wall and observed nuclei protruding into the luminal space, similar to in vivo reports.^56^ This work also showed the capability of in situ modification to existing artificial capillaries. Using two-photon crosslinking, we also show that user-defined obstructions can be added within the microchannels. Although these studies were conducted as proof-of-concept experiments, this strategy can potentially study how user-defined editing relate to local tissue hypoxia, inflammation, and cell death, and whether blockages can be naturally resolved (recanalization).^57^

Since long-term stability of capillaries typically require co-culture with supporting cells,^58^ it is important to show that support cells can be added at any point during active cell culture. We choose fibroblasts-like 10T1/2s as our model support cells due to its wide use in microvascular cocultures.^59^ We designed a new chip with top-open silo arrays (diameter = 150µm) to facilitate seeding of relevant stromal cells during active cell culture. Since the timing of the addition of cells to a microvascular co-culture model has been shown to be critical,^60^ separate seeding ports in our chip provides control over seeding sequence/timing for each cell type during active cell culture. The depth of the silos is optimized such that floor of the silos are ∼50-100µm above the bottom glass interface; this ensures that all interactions between stromal cells (seeded in Ch#2) and artificial capillaries occur within natural 3D collagen matrix and facilitate high-resolution imaging. For PDMS chips, the time from design-to-use takes ∼2 hours which is ideal to rapidly optimize device design. Collagen gelation takes 24hrs while laser-assisted microchannel formation takes few hours; since the fabrication process to sculpt collagen gels via casting and laser cavitation are decoupled from the cell-seeding process, many chips can be fabricated, stored, and used when necessary.

To provide mechanistic biophysical insights related to artificial capillaries, the *in vivo* visualization of transport within the fabricated channels has been augmented with CFD simulations that relate the geometric signature of the artificial capillaries to quantitative, physiologically relevant metrics (namely, flow velocity, wall shear stress, and strain rates) that cannot be, otherwise, fully resolved experimentally, due to the small lumen size range, custom architecture, and scattering due to the collagen matrix. The LES-based gravity-driven simulation scheme was validated with experimental measurements for the straight channel topology and was then representatively applied to the branched, zig-zag, and centrally tapered channel architectures. The numerical estimates (e.g., see **Figs.2, 5,** and **9**) enable prediction of how the in situ architectural edits (in the form of, for example, new branches, occlusions, and tapering) and fibroblast-driven remodeling may redistribute shear and velocity fields, thereby linking structural tunability of the platform to mechanobiological cues that would be locally experienced by endothelial cells.

Our model provides independent control over lumen size (8-35µm), lumen geometry (straight, tapered, curvatures, zig-zag), and associated flow properties, ECM composition (in this case, collagen, 5mg/ml, and HAMA), and seeding location and timing of stromal cells (fibrobasts-like 10T1/2s). We envision artificial capillaries, made using human-derived cells, will be ideally suited to discover new capillary-stroma relationships relevant in tissue development, aging, and diseases which, at present, largely rely upon genetic targeting experiments in mice, which are expensive and time-consuming. This will be important for screening novel therapeutics that target the perivascular niche, particularly relevant considering the recent FDA Modernization Act that allows for alternatives to animal testing. The ability of these chips to be readily shipped to any lab and cultured with clinically relevant cell types and ECM compositions will enable the study of cross-talk between flow, varying based on location, geometry (curvatures) and lumen sizes ECM properties, and perivascular cells necessary to maintain a stable and functional microvascular networks.^61–63^ New avenues of research, such as effects of capillary flow and/or ECM composition on the phenotype and transcriptional profiles of translationally relevant cells,^62, 64–66^ single-cell dynamics of diseased cells,^46^ and capillary obstruction^57^ can be studied. By increasing the size of the silos, larger biologics such as spheroids, organoids or biopsy samples can be reproducibly placed in the vicinity of artificial capillaries. This will have significant impact on various fields such as organ on chip, precise medicine, drug discovery and safety assessment.

## Materials and methods

Please refer to the supplementary information (SI) section. This file also includes additional figures (S1 to S21), movie files (V1 to V16) and associated captions.

## Data availability statement

All data that support the findings of this study are included within the article (and any supplementary files).

## Acknowledgments.

This work was financially supported by funding from the National Institutes of Health, R01 AR083466 and R21 DK136083-01 to PS, and partially supported by SB’s National Science Foundation CAREER AWARD CBET 2339001 (Fluid Dynamics program). We thank Dr Alison Patteson (Physics Department, Syracuse University), for access to Confocal reflectance microscope. We thank Dr. Juan Melero-Martin and Allen Chilun Luo from Harvard Medical School for providing us with ECFCs.

## Materials and Methods

In this work, two chip designs were used. Chip design A was used to generate endothelialized-microchannels embedded in collagen or capillaries-on-a-chip, while Chip design B was used to facilitate co-culture of stromal cells with capillaries-on-a-chip. For the design of each chip, several key steps were involved as described below.

### 1. Chip Design A: Design and fabrication of chips to generate artificial capillaries (SI-1-3)

**Step 1. Design and printing of master molds.** Prepolymer solution was prepared using poly(ethylene glycol) diacrylate (PEGDA, MW= 250Da; Sigma-Aldrich) as the base material, with Phenylbis(2,4,6-trimethylbenzoyl)phosphine oxide (commonly known as Irgacure 819; Sigma-Aldrich) and 2-Isopropylthioxanthone (ITX; Tokyo Chemical Industry) as the photoinitiator and photosensitizer respectively, while 2,2,6,6-tetramethyl-1-piperidinyloxy (TEMPO; Sigma-Aldrich) served as the free radical quencher. A stock solution was prepared by mixing 100 mg of Irgacure 819, 200 mg of ITX, and 4 mg of TEMPO with 40 ml of PEGDA in a centrifuge tube, which was then wrapped in aluminum foil and vortexed to ensure thorough mixing of the chemicals in the solution. This stock solution was then stored at room temperature until use.

Prior to printing, the glass slides were subjected to cleaning in piranha solution (H2SO4 and H2O2; 7:3) with constant stirring at 125 rev/min for 30 minutes, followed by rinsing with ethanol and water and subsequent drying at 65°C in a vacuum oven for an hour. The surfaces of the glass slides were then modified with methacrylate groups by immersing them in a solution containing 3-(trimethoxysilyl)propyl methacrylate (TMSPMA; Sigma-Aldrich) and Toluene (Sigma-Aldrich) (9:1) at 50°C, while maintaining a constant stirring at 125 rev/min. The acrylated glass slides were dried at 65°C in a vacuum oven overnight, and the modified cover slips were affixed to an aluminum print head using a double-sided tape for printing. Custom Digital Light Projection (DLP) platform was designed and built in Soman group.^1^ Briefly, the optical setup consists of a 405 nm CW laser light source (iBEAM SMART 405, Toptica Photonics), DLP development kit (DLP 1080p 9500 UV, Texas Instruments, USA), and a Z-stage (25 mm Compact Motorized Translation Stage, ThorLabs). A rotating diffuser was added to the setup to eliminate laser speckles generated by the Gaussian beam profile, and to convert Gaussian intensity profile into a uniform hat-shaped distribution beam profile before expanding, collimating, and projecting the beam onto the DMD which consists of an array of 1920×1080 micromirrors with a single pixel resolution of around 10µm. Custom MATLAB code was used to slice the models and generate stacks of binary portable network graphics (PNG) image files.

These files were uploaded onto DMD and converted to virtual masks to spatially modulate the laser beam onto a vat of liquid prepolymer solution through an oxygen permeable PDMS window. LabVIEW code was used to control DMD and synchronize it with the upward movement of the print head to photo-polymerize the master mold onto a surface-modified glass coverslip

**Step 2. Fabrication of PDMS microfluidic chips.** PEGDA master molds were used to make the PDMS microfluidic chips using replica casting process. First, polydimethylsiloxane (PDMS, Sylgard 184: Dow Corning Corporation) base and curing agent were thoroughly mixed in a 10:1 mass ratio, degassed under vacuum, and poured onto the master molds. The molds were initially placed in a 52 mm petri dish, then PDMS was poured directly over the micro holes (reverse feature for micro-pillars) and degassed to ensure uniform filling of PDMS in all negative spaces of the mold. PDMS-mold setup was then cured for 10-12 hours at a low temperature (35-40°C) and for an additional 2 hours at a higher temperature (65°C). After cooling, the PDMS layer was gently peeled from the molds and trimmed to size. Holes were punched out of the PDMS to form 3 inlets and 3 outlet ports. Finally, the PDMS was irreversibly bonded to a glass coverslip (0.15mm thick) using oxygen plasma treatment (60-seconds exposure).

**Step 3. Collagen gelation within central chamber (Ch#2) of PDMS chips.** The microfluidic chip surfaces were modified with (3-aminopropyl) triethoxysilane (APTES) and glutaraldehyde (GA) to immobilize the collagen and prevent detachment from chamber surfaces. First, chips were sequentially immersed in a solution of (i) 10% (APTES, Sigma-Aldrich) in ethanol for 1 hour, followed by three ethanol rinses, and (ii) 2.5% glutaraldehyde (GA, Sigma-Aldrich) solution in deionized water for 1 hour, followed by three rinses using deionized water, and (iii) sterilization under UV light overnight. Type I collagen solutions at varying concentrations (Rat tail tendon, ibidi, Germany) were prepared using established protocols. Briefly, stock solution of collagen (10 mg/ml) was diluted to 5 mg/ml with 1x medium (DMEM, ThermoFisher) and 10x medium (HBSS, ThermoFisher) such that the final salt concentration is 1x and it is neutralized to pH 7.2–7.4 by adding calculated amount of 1M NaOH. All the reagents, media and collagen were placed in ice-bath throughout the neutralization process. Final pH was measured using pH strips. Collagen solution was pipetted into Ch#2 of the chips and incubated at 37°C for 20 minutes to allow gelation of the matrix. Then, 1x PBS supplemented with 1% Penicillin-Streptomycin (10,000 U/mL, Thermo Fisher Scientific) was pipetted into Ch#1 and Ch#2 to prevent crosslinked collagen from dehydration and bacterial contamination. The silos were visualized under reflectance microscopy, and then the chips were stored in an incubator for 24 hours to ensure robust crosslinking of collagen and consistent channel sizes during laser scanning experiments. All femtosecond laser scanning experiments were performed after 24hours at user-defined time-points.

**Step 4. Microchannel generation using femtosecond laser assisted cavitation.** Custom-built fs-laser setup was designed and built by combining a Ti:Sapphire fs laser (Coherent, Chameleon, USA) with a Zeiss Microscope (Observer Z1, Germany). In this setup, an 800 nm femtosecond laser beam with a repetition rate of 80 MHz was focused inside crosslinked collagen within the microfluidic chips using a water immersion objective (40x, NA=0.8, Leica). The microscope stage, controlled using Visual Basic script that was integrated with Axio Vision software (Zeiss, Germany), was used to create microchannels of different architecture and dimensions. The laser dosage was changed by modulating the average power of the laser using a polarization-based power-tuning system or by modulating the scanning speed of the stage.

Multiple scans were performed to achieve the desired lumen sizes of the channels and they were visualized under brightfield or reflectance confocal microscopy.

### 2. Chip Design B. Design and fabrication of chips to enable co-culture of stromal cells with artificial capillaries-on-chips. (SI-12 to SI-17)

**Step 1. Design and printing of master molds.** Master mold for design B required an assembly of a (1) bottom-part (negative replica of three-microchambers with two posts, **SI-12**), and (2) top-part with a conical plug with two holes **(SI-13)** Both top and bottom parts were printed as described earlier and assembled. **(SI-14)**

**Step 2. Fabrication of PDMS microfluidic chips.** Replica casting, as described for chip design A, was used to make the PDMS chip. **(SI-15, V-8)**

**Step 3. Collagen gelation within central chamber (Ch#2) of PDMS chips.** Plug with two pins was printed **(SI-16)** and assembled into the central chamber of the PDMS chip. **(SI-17, V-9)** Then, process described earlier was used to gel collagen around and under the two-pins (diameter = 150μm). The gap between the inserted pin bottom and the glass surface is ∼50-100µm. Post-gelation, the plug was removed leaving behind two silos of diameter 150μm inside collagen in Ch#2 of the chips. Then, 1x PBS supplemented with 1% Penicillin-Streptomycin was pipetted into Ch#1 and Ch#3 and into silos to prevent crosslinked collagen from dehydration and bacterial contamination. Trapped air bubbles, if observed above the silos, were removed by aspirating them with a syringe.

**Step 4. Microchannel generation using femtosecond laser assisted cavitation.** As described earlier, 24 hours after step 3, two microchannels of defined lumen sizes were made within collagen on either side of the silos. Note that the microchannels are made ∼50-100µm inside collagen above the bottom glass coverslips.

### 3. Cell culture

In this study, three types of cells were used: Human Umbilical Vein Endothelial Cells (HUVECs), Human Endothelial Colony-Forming Cells (ECFCs), and fibroblast-like 10T1/2 cells. HUVECs and 10T1/2 cells were purchased from American Type Culture Collection (ATCC, Manassas, VA) and ECFCs were provided by Allen Chilun Luo (Boston children hospital).

HUVECs were cultured using EGM-2 Endothelial Cell Growth Medium-2 BulletKit (Lonza) containing EBM-2 Basal Medium (CC-3156) and EGM-2 SingleQuots Supplements (CC-4176) required for growth of Endothelial Cells. 1% penicillin/streptomycin (Thermo-Fisher Scientific) was added as antibiotics. This culture system contains 2% Fetal Bovine Serum (FBS). ECFCs were culture in same media but with total of 20% FBS. 10T1/2 cells, derived from a C3H mouse embryo cell were cultured in a cell growth media consisting of a Basal Medium Eagle (BME, GIBCO#21010-046) supplemented with 10% FBS, 1% penicillin/streptomycin and 1% GlutaMAX (GIBCO#35050-061). All the cells were cultured in T75 vented cell culture flasks in a standard incubator at 37°C with 5% CO2, with media changes occurring every 2-3 days. Cells were maintained at 37 °C in a humidified incubator with 5% CO₂ and subcultured at 80–90% confluency using 0.25% trypsin-EDTA. Cells between passages 3 and 6 were used for all experiments. Upon reaching 80%-90% confluency, the cells underwent a single wash with 1 mL of 1xPBS, followed by treatment with 3 mL 1x TrypLE Select (TrypLE; Thermo Fisher Scientific, catalog no. 12563029) incubated at 37°C with 5% CO_2_ for 3 minutes. The reaction was neutralized by adding 5 mL of respective growth medium and centrifuging at room temperature at 150G for 5 minutes.

### 4. Cell seeding and setup to achieve unidirectional perfusion through microchannels

To achieve endothelialization of the laser cavitation-assisted microchannels, 20 µL of HUVEC cell solution (1 or 0.5 × 10⁶ cells/mL) was manually pipetted into one of the side channels (Ch#1 or Ch#3), allowing cells to migrate into the adjacent microchannels in Ch#2. To facilitate cell entry into the microchannels, the device was initially tilted at a 90° for 10 minutes. Then, the opposite side channel was also filled with cell suspension and held at a 90° for another 10 minutes to enhance uniform seeding. Following seeding, all four inlets were filled with endothelial growth medium, and the devices were incubated in 35 mm cell culture dishes under static conditions (37 °C, 5% CO₂, and 90% humidity) for 24 hours prior to initiating perfusion culture. For experiments with Chip design B, after ensuring the microchannels are fully lined with HUVECs, model stromal cells (fibroblasts-like 10T1/2s), total of 10 µL of cell suspension (1 × 10⁶ cells/mL) was seeded in the silos in collagen wall through the funnel shaped openings (reservoir) in the PDMS chips, and media in the central silo inlets were aspirated out using 20 µL pipette. Both monocultured chips (design A) and co-culture chips (design B), we cultured using Endothelial Cell Growth Medium with high serum content. To maintain a simple experimental setup, we adopted a pseudo-perfusion approach by attaching 1 mL pipette tips to the inlets of one of the side chambers and filling them with specific volume of media, which corresponded to defined pressure head condition. The fluid velocity in the microchannels was estimated by tracking the movement of microbeads. Briefly, a total of six microchannels in collagen wall in Ch#2 (lumen size=40μm) were generated using laser-assisted cavitation. Two 1 mL pipette tips were trimmed at the ends to fit into the reservoirs of one of the side channels (either Ch#1 or Ch#3), while the opposite channel was left open to allow the fluid to drain away. A calculated volume of PBS containing a mixture of 0.5 μm (Polybead, Polysciences) and 6 μm beads (Spherotech) was added to the pipette tips. Both tips were filled with the same volume of PBS, and the bead flow into Ch#3 was recorded using a high-speed camera (Hamamatsu, Japan) at 40 frames per second. The movement of 6 μm beads exiting into the downstream channel was tracked using FIJI, and corresponding velocities were calculated. Velocity measurements were averaged across five individual beads in each of the 4 different channels. Simulation was carried out to expand the experimental results and assess flow parameters within the microchannels; see next.

### 5. Experimentally validated computational fluid dynamics (CFD) modeling (SI-4)

High-fidelity Large Eddy Simulation (LES) scheme on ANSYS Fluent 2024 R1 was used to develop gravity-driven flow conditions, with the kinetic energy transport model resolving fluctuations at the subgrid scales. We reconstructed the channel geometries using ANSYS Workbench 2024 R1, and to mesh the geometries ICEM CFD 2024 R1 was utilized. To ensure proper domain preparation for numerical simulation, a grid refinement study was conducted. The straight microchannel (diameter 40 µm) was discretized using seven levels of spatially graded, unstructured tetrahedral mesh elements, ranging from approximately (0.017, 0.031, 0.049, 0.073, 0.096, 1.5 and 1.7 million respectively). The sensitivity analysis evaluated variables such as the flow resistance R (expressed in Pa·min/mL) to the simulated fluid flow, calculated as 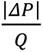, where |Δ𝑃| in Pa represents the inlet-to-outlet pressure gradient driving the flow and Q denotes the volumetric flux in mL/min. To determine the final mesh configuration, we have performed variance calculations using all seven mesh densities, as well as separately for the three finest meshes (0.096, 1.5, and 1.7 million elements). The variance values are:

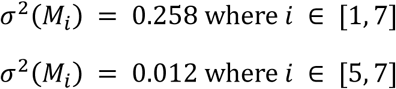

Based on the optimal balance of computational efficiency and accuracy observed in the simulations, the 1.7 million-cell mesh resolution was chosen for the comprehensive analysis. For the other complex and branched geometries, the mesh was generated in accordance with this chosen resolution. This selection aligns with comprehensive analyses of grid convergence and computational stability conducted for fluid flow simulations within microchannels under five distinct pressure head conditions. To evaluate the numerical stability of the simulations, the Courant–Friedrichs–Lewy (CFL) number was computed. In this study, the CFL number was found to be 0.885, and since the condition (CFL < 1) is satisfied, the numerical scheme is considered stable throughout the simulation.

The fluid flow within the microchannels was replicated for five different volumes (1 mL, 0.5 mL, 0.25 mL, 0.10 mL and 0.05 mL). We used pressure inlet and pressure outlet boundary conditions (pressure-driven flow). A no-slip boundary condition (zero velocity) was applied at the channel walls, and the pressure-driven simulations were conducted using a time step of 0.0001 seconds. For the flow solution time of 0.10 s, we chose the total number of time steps = 1000. The simulations were performed using a segregated solver incorporating pressure–velocity coupling and second-order upwind spatial discretization. For the transient formulation, a bounded second-order implicit scheme was employed to ensure numerical stability and prevent non-physical oscillations. Convergence was assessed through minimizing the mass continuity and velocity component residuals. The orders-of-magnitude for the continuity residuals were *O*(10^-4^) in the straight channel, *O*(10^-4^) in the zig-zag channel, *O*(10^-3^) in the branched channel, and *O*(10^-4^) in the mid-tapered channel. Likewise, at convergence, the orders-of-magnitude for the velocity component residuals were *O*(10^-7^) in the straight channel, *O*(10^-7^) in the zig-zag channel, *O*(10^-6^) in the branched channel, and *O*(10^-6^) in the mid-tapered channel. The simulations generated pressure-driven flow, with the solver set to transient. Considering water as a working fluid passing through the microchannels, the water density (ρ) was set at 998.2 kg/m^3^ in the simulations, with 0.001003 m^2^/s as kinematic viscosity (𝜈). For 1 mL, the inlet pressure was set at 519 Pa, and the outlet pressure was defined at 0 Pa. Similarly, the other four simulations were conducted using the inlet pressure values of 332.94 Pa, 215.43 Pa, 127.30 Pa and 78.34 Pa corresponding to the respective volumes for the straight microchannel (length = 800 µm, diameter = 40 µm). A similar simulation procedure was applied to the complex and branched capillary geometries to predict the distribution of velocity and wall shear stress throughout the domains (note that for the complex domains, only 519 Pa was used).

To replicate the experimentally measured velocities within the numerical domain of the straight channel, the standard deviation values associated with five distinct pressure heads were incorporated. The experimental data indicated bead velocities of 17.12, 9.0, 7.6, 3.8, and 2.4 mm/s, corresponding to pressure heads of 519, 332.94, 215.43, 127.30, and 78.34 Pa, respectively. The respective standard deviations for these velocities were 0.53, 0.36, 0.30, 0.21, and 0.27.

By utilizing the Hagen-Poiseuille’s equation for the laminar flow of an incompressible, Newtonian fluid through a circular microchannel, we determined the annular region to obtain the predicted velocities for all the pressures. The Hagen-Poiseuille’s equation for velocities as a function of radial distance is

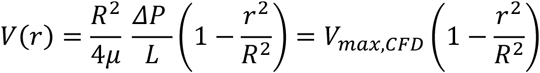

In the above equation, *R* = radius of the straight microchannel = 20 µm, *ΔP* = corresponding pressure difference, *L* = length of the microchannel = 800 µm, µ = dynamic viscosity. 𝑉*_max,CFD_* was used from the simulation data for distinct pressure heads to select the annular regions. Based on the standard deviation for 519 Pa, we obtained *V*(r_1_) and *V*(r_2_) from 17.12 ± 0.53 mm/s which implied that [𝑟_1_, 𝑟_2_] = [16.59, 17.65] mm/s. We obtained the numerical average velocity for the annular region, where three grid points were analyzed, was 17.82 mm/s at 519 Pa. Performing the similar procedure, we calculated the numerical velocities for all the pressure heads, and the results are plotted on the bar diagram. The wall shear stress across the entire domain is represented, and Fig. 2 includes the line plot of area weighted average of wall shear stress corresponding to five distinct volumes for the simple circular channel (approximately 6.45, 4.15, 2.69, 1.59, 0.98 Pa for volumes in the order of higher to lower, respectively). The associated annular region’s velocities are plotted where the red marker defines the experimental results for each of the volumes, respectively. Additionally, we predicted the streamwise velocity profile and wall shear stress throughout the entire domain for complex and branched capillaries. Three additional distinct intricate microchannels were numerically studied for 519 Pa pressure-gradient; see Figs. 5 and 9. The number of unstructured tetrahedral elements to discretize the complex channel domains was selected based on the mesh specifications in our simpler domain (straight microchannel) following the relation:

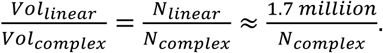

### 6. Permeability characterization of artificial capillaries (SI-6)

A fluorescein isothiocyanate (FITC)-conjugated dextran of 70ka and 4kDa (Sigma) was used to test the barrier function of endothelialized chips. Chips without HUVECs were used as controls. HUVEC-lined chips were prepared as previously described for 3 days to allow for healthy monolayer formation. Then FITC-dextran solution in endothelial growth medium was prepared and introduced into one side chamber of the chip at concentration of 10 μg/mL, while the opposite side chamber was filled with dextran-free medium. The diffusion of FITC-dextran along and across the microchannels in Ch#2 were monitored using timelapse images taken at different time intervals (0, 5, 15, 30, 45, 60, 120, and 180 min) using fluorescence microscope (excitation/emission: 495/519 nm). To assess diffusion and calculate permeability across a microchannel interface, time-lapse fluorescence images were acquired and saved as a TIFF stack. Since we have large stacks of images, we downsized the file by taking every 10^th^ slices. Fluorescence intensity profiles along a specified line (ROI) were extracted from every 10th image slice to track how fluorescence progressed into the gel over time. For each sampled time point, the maximum fluorescence intensity along the line was recorded and analyzed using FIJI image processing software. The maximum values were then plotted as a function of time of diffusion. The initial slope of this curve was computed using a linear fit over the first five data points. Permeability was then estimated using the equation 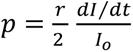; where r is the channel radius (15 µm), and 𝑑*I*⁄𝑑*t* is the initial rate of increase in fluorescence intensity, and *I_o_* is the initial source intensity at the interface. Experiments were performed in triplicates,

### 7. Cell morphology and immunostaining

To visualize HUVEC morphology within microchannels, chips were fluorescently stained and imaged using an LSM 980 confocal microscope (Zeiss, Germany). The samples were fixed using a 4% formaldehyde (Sigma-Aldrich) was pipetted in Ch#1 and #3 for 30 minutes at room temperature and washed three times with 1x PBS. Cells within Ch#2 of the device were permeabilized with 0.5% TritonX-100 in 1x PBS for 30 minutes and again washed 3 times with PBS. For immunostaining, samples were blocked with 1% Bovine serum Albumin (BSA; A3160801, Thermo Fisher) for 1 hour at room temperature, followed by thorough washing with 1x PBS. To ensure complete removal of residual serum, 1 mL pipette tips were inserted into two inlets of one side channel and filled with PBS to generate a pressure differential, effectively flushing the remaining serum into the opposite side channel which was washed again with PBS. Chips were then incubated with primary antibodies overnight at 4°C. After incubation, chips were washed 3x with PBS as previously described, followed by incubation with secondary antibodies. Primary antibodies included CD31 Recombinant Rabbit Monoclonal Antibody (clone 078; #MA529475, Thermo Fisher) and CD44 Monoclonal Antibody (clone IM7; #14044182, Thermo Fisher), both used at a 1:200 dilution in blocking buffer containing 0.2% BSA, 0.1% Tween-20, and 0.3% Triton X-100 (BTT). Secondary antibodies included Goat anti-Rabbit IgG (H+L), Alexa Fluor 488 conjugate (#A11034, Thermo Fisher), and Donkey anti-Rat IgG (H+L), Alexa Fluor 568 conjugate (#A78946, Thermo Fisher), each diluted 1:1000 in BTT. All samples were stained with DAPI (4′,6-diamidino-2-phenylindole; #62248, Thermo Fisher) at a dilution of 1:1000 for 5 minutes at room temperature to visualize cell nuclei. For f-actin visualization, selected chips were incubated with Alexa Fluor 680 Phalloidin (#A22286, Thermo Fisher) diluted 1:250 in 10% horse serum for 45 minutes at room temperature. After staining, samples were thoroughly washed and stored in 35 mm petri dishes, covered with aluminum foil to protect from light and prevent photobleaching of fluorophores.

### 8. Image and data processing

For both chip designs (A and B), z-stacked fluorescence images were acquired, and processed using FIJI (ImageJ). Maximum intensity projections were generated to visualize overall cell morphology. For better visualization of lumen structure and endothelial coverage, z-stacks were resliced orthogonally along the channel axis. The 3D Viewer plugin was used to generate volumetric renderings and capture high-resolution snapshots to illustrate lumen architecture and cellular coverage within the lumens. Endothelial thickness was estimated from CD31-immunostained microchannels by analyzing a single representative confocal image take from 3 independent chips. **(SI-8)** Image taken from the center of lumens were selected to provide a consistent cross-sectional view to determine the . A median filter was applied to reduce noise and enhance image quality. Intensity profiles were extracted perpendicular to the endothelial lining, and the full width at half maximum (FWHM) of each fluorescence peak was used to approximate the local endothelial thickness.

Nuclear orientation was analyzed by fitting ellipses to individual nuclei and measuring the angle between the major axis of each nucleus and the longitudinal axis of the channel. All measurements were performed on at least three independent regions per sample, and results were averaged to obtain representative values. Briefly, nuclei were automatically counted from z-stacked image volumes using a custom MATLAB script. TIFF stacks were loaded slice by slice into a 3D array, followed by 3D Gaussian smoothing (σ = 1) to reduce noise. Otsu’s method (MATLAB) was used to determine a global threshold for binarization. Small objects below a minimum voxel size (20 voxels) were removed to eliminate background noise. The cleaned binary volume was then processed using 26-connectivity to identify and label distinct 3D connected components, corresponding to individual nuclei. The total number of nuclei was obtained by counting the number of connected components in the volume. Quantitative analyses and data visualization were performed using MATLAB (R2023a). Nuclear orientation data, obtained from image processing, were visualized using a custom-generated rose diagram. The orientation data, ranging from 0° to 180°, were plotted as a normalized polar histogram using 18 bins (10° resolution). Histogram normalization was performed by converting the frequency counts into probabilities, enabling comparative visualization independent of sample size. The maximum probability value was scaled to 0.2 to ensure consistent plot scaling and emphasize relative distribution differences. Polar plot was generated to depict the distribution of nuclear alignment relative to the longitudinal axis of the channel. To estimate the area occupied by each cell within endothelialized microchannels, z-stacked 3D confocal images were processed using custom MATLAB scripts. Image stacks were loaded into a 3D array, smoothed with a 3D Gaussian filter (σ = 1), and segmented using Otsu’s thresholding. Small objects were removed, and connected components were identified using 26-connectivity.

The number of nuclei per channel was quantified as the total number of 3D connected components. Assuming a cylindrical lumen geometry, the inner surface area of each lumen was calculated using 𝐴 = 2𝜋𝑟ℎ, where 𝑟 is the measured lumen radius and ℎ is the lumen length. Using 3D segmentation and connected component analysis, we quantified the number of nuclei present in each capillary lumen and calculated the average size of each cell. Based on the calculated surface area of the cylindrical lumens, the average area per cell varied between 1024 µm² and 1433 µm². The average area per cell was obtained by dividing the surface area by the number of nuclei. To evaluate how cellular morphology varies with lumen size, area-per-cell values were compared across channels with diameters using scatter plots, box plots, and one-way ANOVA with Tukey’s post-hoc tests.

Our analysis using one-way ANOVA revealed a statistically significant effect of lumen diameter on cell spreading behavior, as measured by average area per cell (F(3,8) = 5.46, p = 0.024). Post-hoc Tukey’s test indicated a significant difference between the 28 µm and 40 µm lumen groups (p = 0.018), with cells in smaller lumens exhibiting larger spreading areas. Together, these analyses provide quantitative evidence that lumen diameter significantly influences cell coverage and spatial organization in engineered capillary models.

| Sample | Lumen Diameter | Num Cells | Total Area | Area PerCell |
| --- | --- | --- | --- | --- |
| Ch1 | 10 | 1 | 1570 | 1570 |
| Ch2 | 10 | 1 | 1479.289 | 1479.289 |
| Ch3 | 10 | 1 | 1570 | 1570 |
| Ch4 | 10 | 1 | 1413 | 1413 |
| Ch5 | 10 | 1 | 1465.333 | 1465.333 |
| Ch6 | 10 | 1 | 1664.2 | 1664.2 |
| Ch7 | 25 | 1 | 1242.603 | 1242.603 |
| Ch8 | 25 | 1 | 1433.772 | 1433.772 |
| Ch9 | 25 | 1 | 1331.36 | 1331.36 |
| Ch10 | 25 | 1 | 1380.67 | 1380.67 |
| Ch11 | 25 | 1 | 1491.123 | 1491.123 |
| Ch12 | 25 | 1 | 1202.519 | 1202.519 |
| Ch13 | 30 | 1 | 1174.729 | 1174.729 |
| Ch14 | 30 | 1 | 1193.2 | 1193.2 |
| Ch15 | 30 | 1 | 1248.15 | 1248.15 |
| Ch16 | 30 | 1 | 1256 | 1256 |
| Ch17 | 30 | 1 | 1198.909 | 1198.909 |
| Ch18 | 30 | 1 | 1205.76 | 1205.76 |
| Ch19 | 35 | 1 | 1164.94 | 1164.94 |
| Ch20 | 35 | 1 | 1092.894 | 1092.894 |
| Ch21 | 35 | 1 | 1194.81 | 1194.81 |
| Ch22 | 35 | 1 | 1164.94 | 1164.94 |
| Ch23 | 35 | 1 | 1226.253 | 1226.253 |
| Ch24 | 35 | 1 | 1156.842 | 1156.842 |
| Ch25 | 40 | 1 | 1024.123 | 1024.123 |
| Ch26 | 40 | 1 | 1064.61 | 1064.61 |
| Ch27 | 40 | 1 | 1109.467 | 1109.467 |
| Ch28 | 40 | 1 | 1086.824 | 1086.824 |
| Ch29 | 40 | 1 | 1046.667 | 1046.667 |
| Ch30 | 40 | 1 | 1116.444 | 1116.444 |

One-Way ANOVA Results
| Source | SS | df | MS | F | Prob>F |
| --- | --- | --- | --- | --- | --- |
| Groups | 743656.812 | 4 | 185914.203 | 37.28039319 | 3.40388E-10 |
| Error | 124672.9092 | 25 | 4986.916368 |  |  |
| Total | 868329.7212 | 29 |  |  |  |

Group-wise Descriptive Statistics
| LumenDiameter | Mean_AreaPerCell | SD | SE |
| --- | --- | --- | --- |
| 10 | 1526.970333 | 91.21073745 | 37.23662764 |
| 25 | 1347.007833 | 110.8626089 | 45.25947054 |
| 30 | 1212.791333 | 32.22404249 | 13.15541026 |
| 35 | 1166.779833 | 44.46885352 | 18.15433343 |
| 40 | 1074.689167 | 36.17730398 | 14.7693225 |

Tukey HSD Post-hoc Comparisons
| Group1 | Group2 | Lower CI | Mean Diff | Upper CI | P Value |
| --- | --- | --- | --- | --- | --- |
| 1 | 2 | 60.2222001 | 179.9625 | 299.7027999 | 0.001465875 |
| 1 | 3 | 194.4387001 | 314.179 | 433.9192999 | 4.4779E-07 |
| 1 | 4 | 240.4502001 | 360.1905 | 479.9307999 | 4.50815E-08 |
| 1 | 5 | 332.5408668 | 452.2811667 | 572.0214666 | 1.02964E-08 |
| 2 | 3 | 14.4762001 | 134.2165 | 253.9567999 | 0.022573859 |
| 2 | 4 | 60.4877001 | 180.228 | 299.9682999 | 0.00144201 |
| 2 | 5 | 152.5783668 | 272.3186667 | 392.0589666 | 5.00058E-06 |
| 3 | 4 | -73.7287999 | 46.0115 | 165.7517999 | 0.790107636 |
| 3 | 5 | 18.36186677 | 138.1021667 | 257.8424666 | 0.018093838 |
| 4 | 5 | -27.6496332 | 92.09066667 | 211.8309666 | 0.191770221 |

### 9. Generation of defined occlusion within artificial capillaries (SI-11)

To create an occlusion in microchannels, a 2.5% (w/v) GelMA solution mixed with 0.25% (w/v) photoinitiator was injected into the microchannels through channel #1 or #3. The GelMA solution infiltrated the microchannels and diffused into the surrounding collagen matrix, as later confirmed by crosslinking observed beyond the channel walls. Using two-photon polymerization (2PP) (laser power 0.3W, objective (20X, NA=0.22, Zeiss), we selectively crosslink the GelMA within the microchannels. Point scanning was performed at a rate of 100 µm/s. A log-pile architecture was fabricated, consisting of orthogonally stacked filaments with a 1 µm gap in both the lateral (X) and axial (Z) directions. Scanning strategies were applied such that crosslinking starts a little away from the microchannel and invades into the microchannel. Since we are using point scanning to crosslink GelMA, we wanted to ensure that the crosslinked GelMA remains anchored to the microchannel wall with its parts that were crosslinked outside the wall in the bulk collagen.

### 10. Capillary-sized microchannel fabrication on a chip using HAMA (SI-20)

Lyophilized hyaluronic acid methacrylate (HAMA; cat# HA-Methacrylate-100k, HAworks) was dissolved in phosphate-buffered saline (PBS) to achieve a final concentration of 4 mg/mL. Lithium phenyl-2,4,6-trimethylbenzoylphosphinate (LAP; Sigma-Aldrich) was used as the photoinitiator to enable the photopolymerization of HAMA via ultraviolet (UV) illumination. The HAMA solution was introduced into chamber 2 and crosslinked under UV light using a UVC-LCB-010 UV box (B9Creations) for 10 seconds. Before laser cavitation of crosslinked HAMA, side chambers of the chip will filled with PBS supplemented with antibiotics. Microchannels of varying lumen diameters (3-40 µm) were fabricated and perfused with fluorescein isothiocyanate (FITC)-Dextran (70 kDa; Sigma Aldrich) into one side chamber, followed by imaging with a LSM 980 confocal microscope (Zeiss, Germany).

---

**SI-1.**
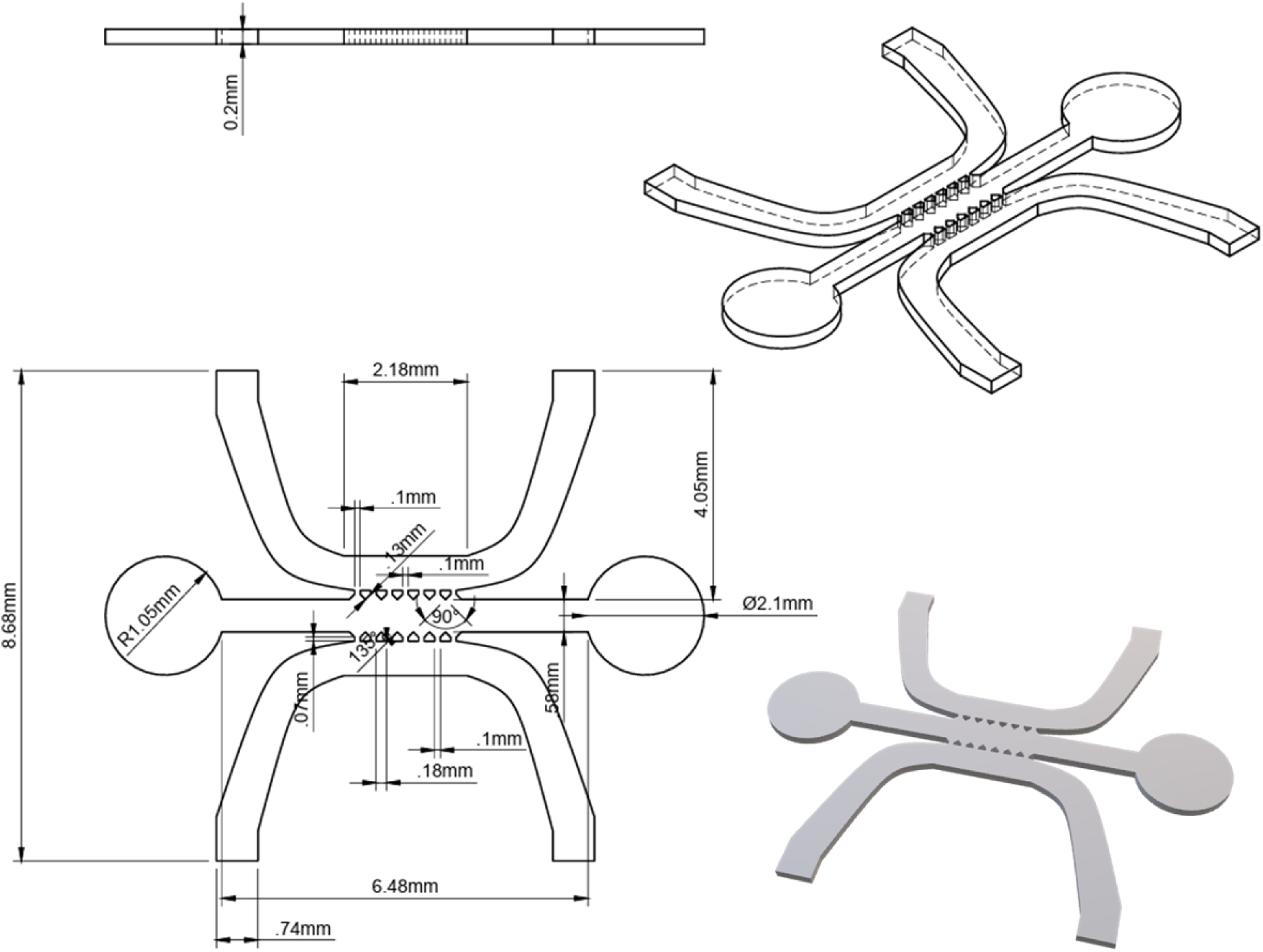
CAD file used to 3D print master molds for chip design A

**SI-2.**
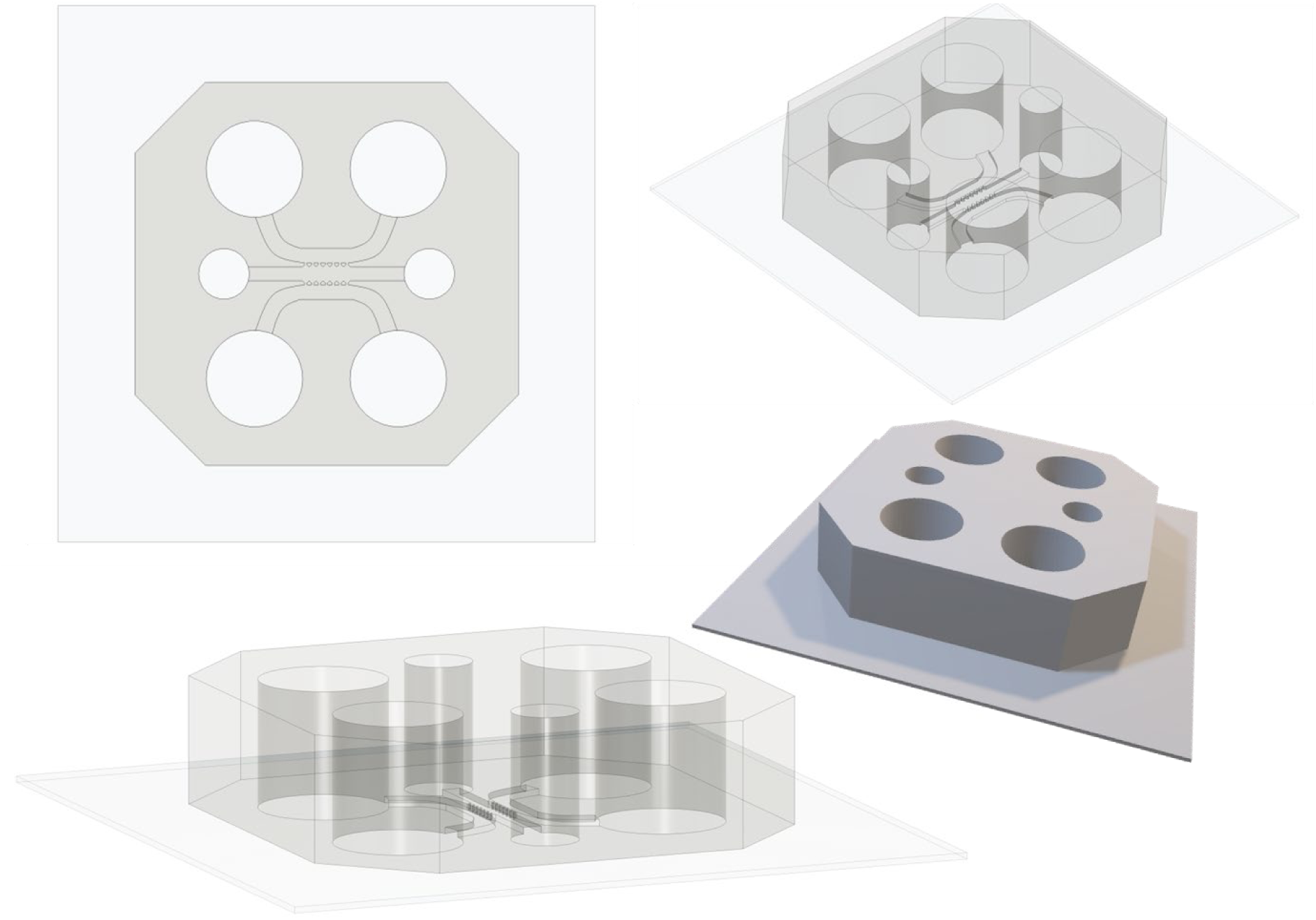
3D CADs showing the final PDMS chip using design A.

**SI-3.**
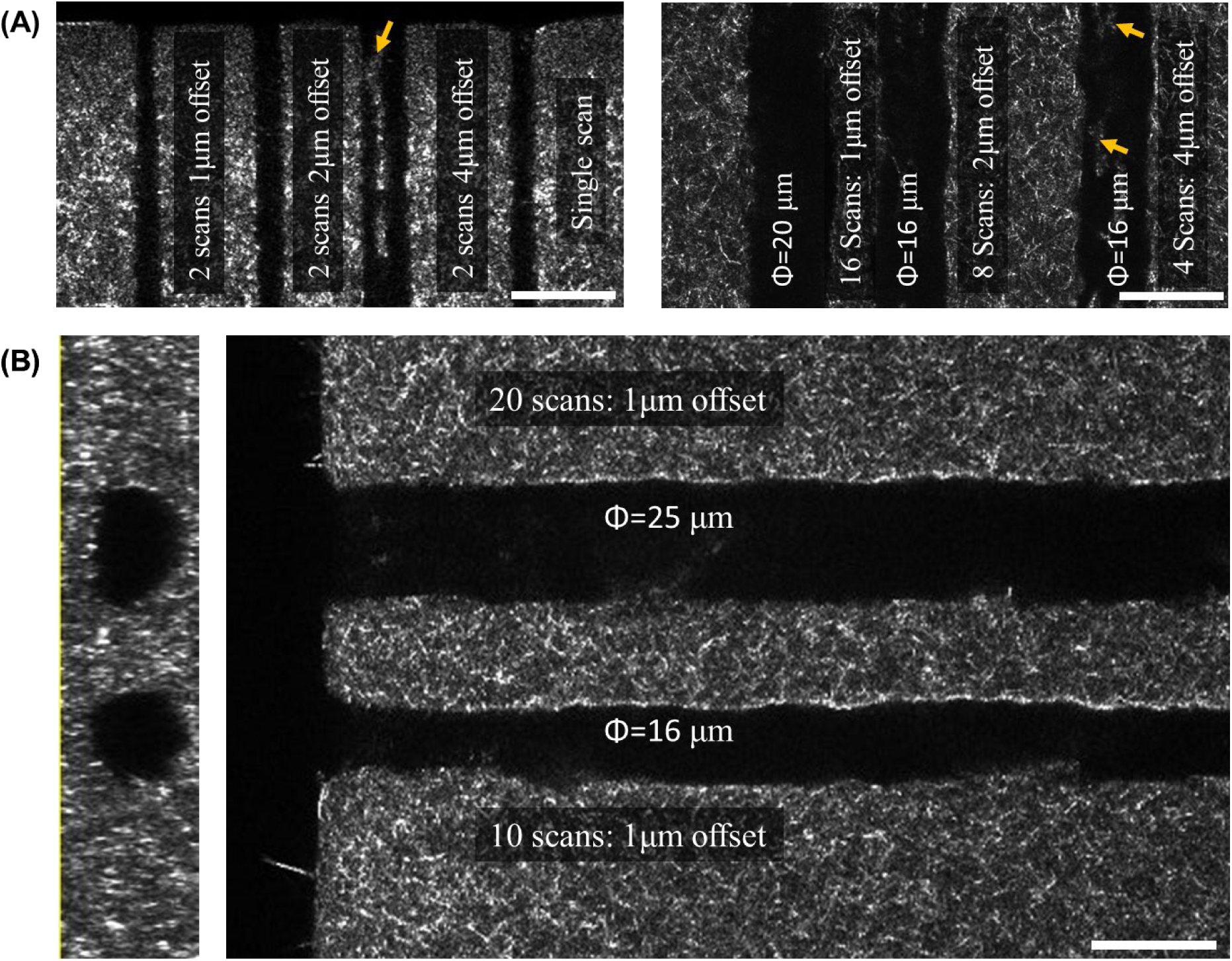
(A-B) Confocal reflectance microscopy top and cross-sectional images showing the effects of varying lateral offsets between fs-laser scanning in collagen. Arrows point to defects when offset was greater than 1µm. Scale bar: 25 μm.

**SI-4.**
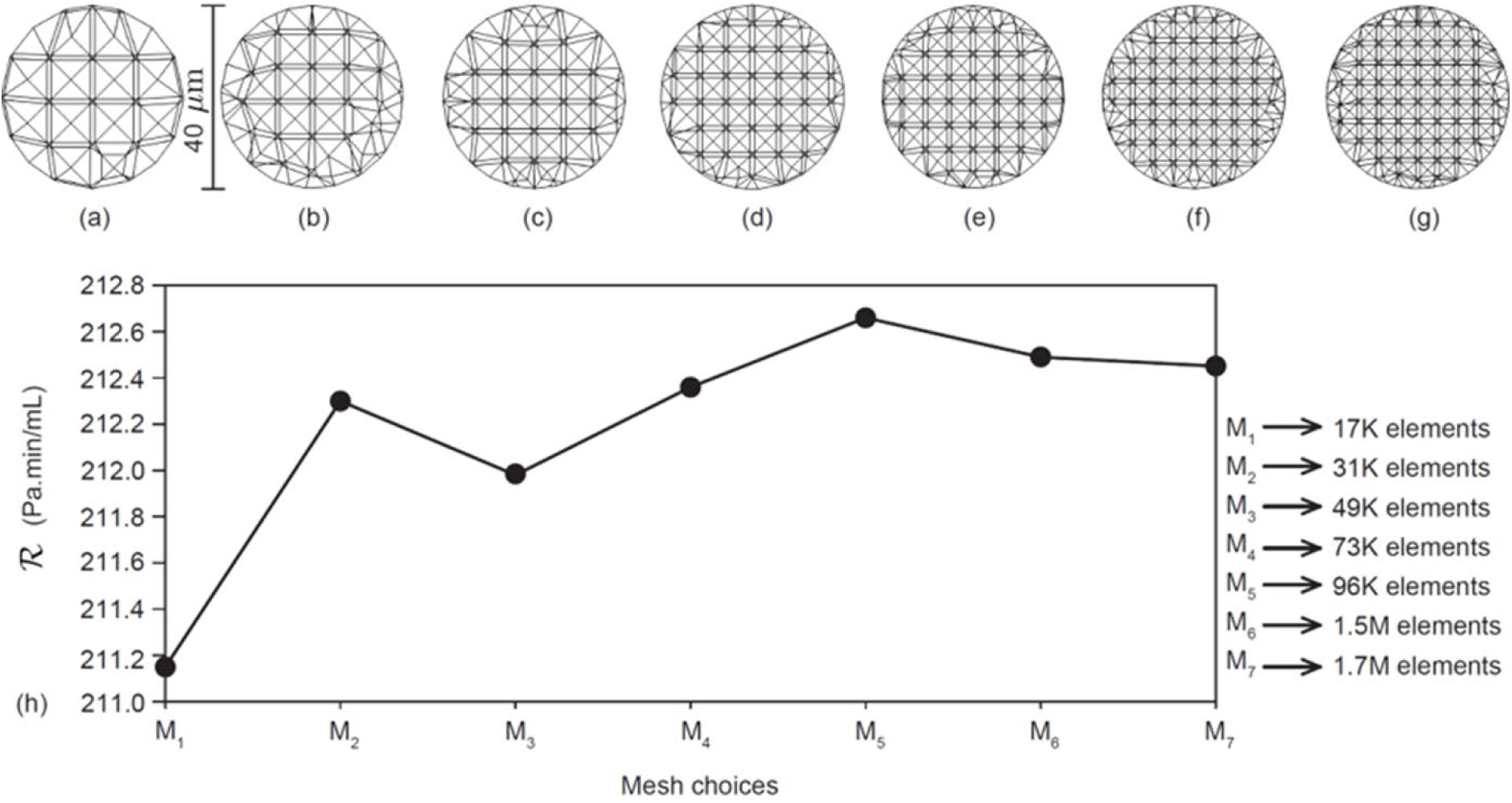
Grid sensitivity analysis: Panels a-g respectively show the mid-section in the linear channel, for the meshed domains with 0.017, 0.031, 0.049, 0.073, 0.096, 1.50 and 1.70 million graded, unstructured tetrahedral elements, respectively. Panel h plots the flow resistance outcome from simulations performed in each mesh. The resistance is calculated as |𝛥𝑃|/𝑄, where |Δ𝑃 | in Pa represents the inlet-to-outlet pressure gradient driving the microchannel flow and Q denotes the volumetric flux in mL/min. To determine the final mesh configuration, we have performed variance calculations (on the flow resistance values) using all seven mesh densities, as well as separately for the three finest meshes (0.096, 1.5, and 1.7 million elements). The variance values were 0.258 when all meshes were considered and 0.012 for the last three meshes. Accordingly, the finest mesh from the last three, i.e., the one with 1.7 million tetrahedral elements 1.7 was selected as a numerical domain for the comprehensive analysis.

**SI-5.**
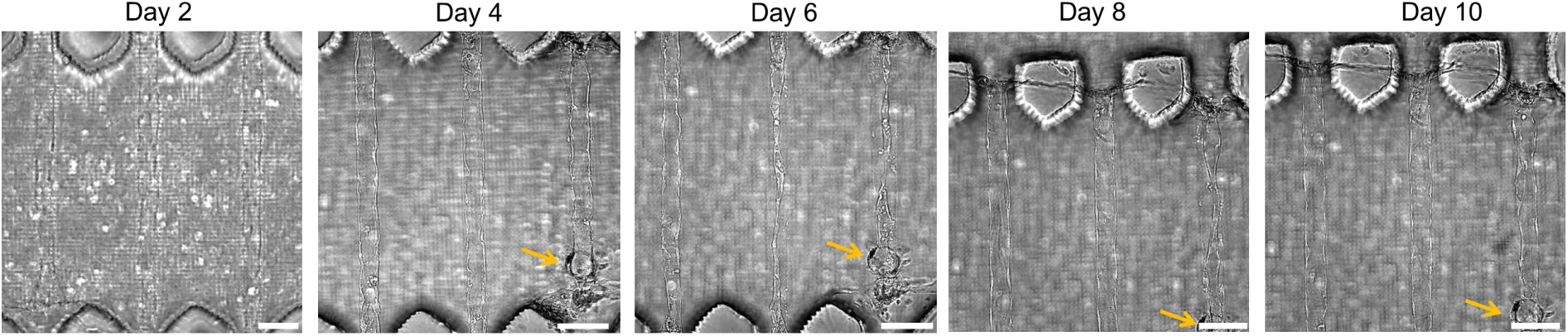
Brightfield images of microchannels after HUVEC seeding at 1M/ml concentration. Arrow indicates a blockage formed in one of the channels, which can be removed by simply reversing the media flow. For seeding concentrations of 0.5M/ml or less, no blockages were detected. Scale bar: 100 μm

**SI-6.**
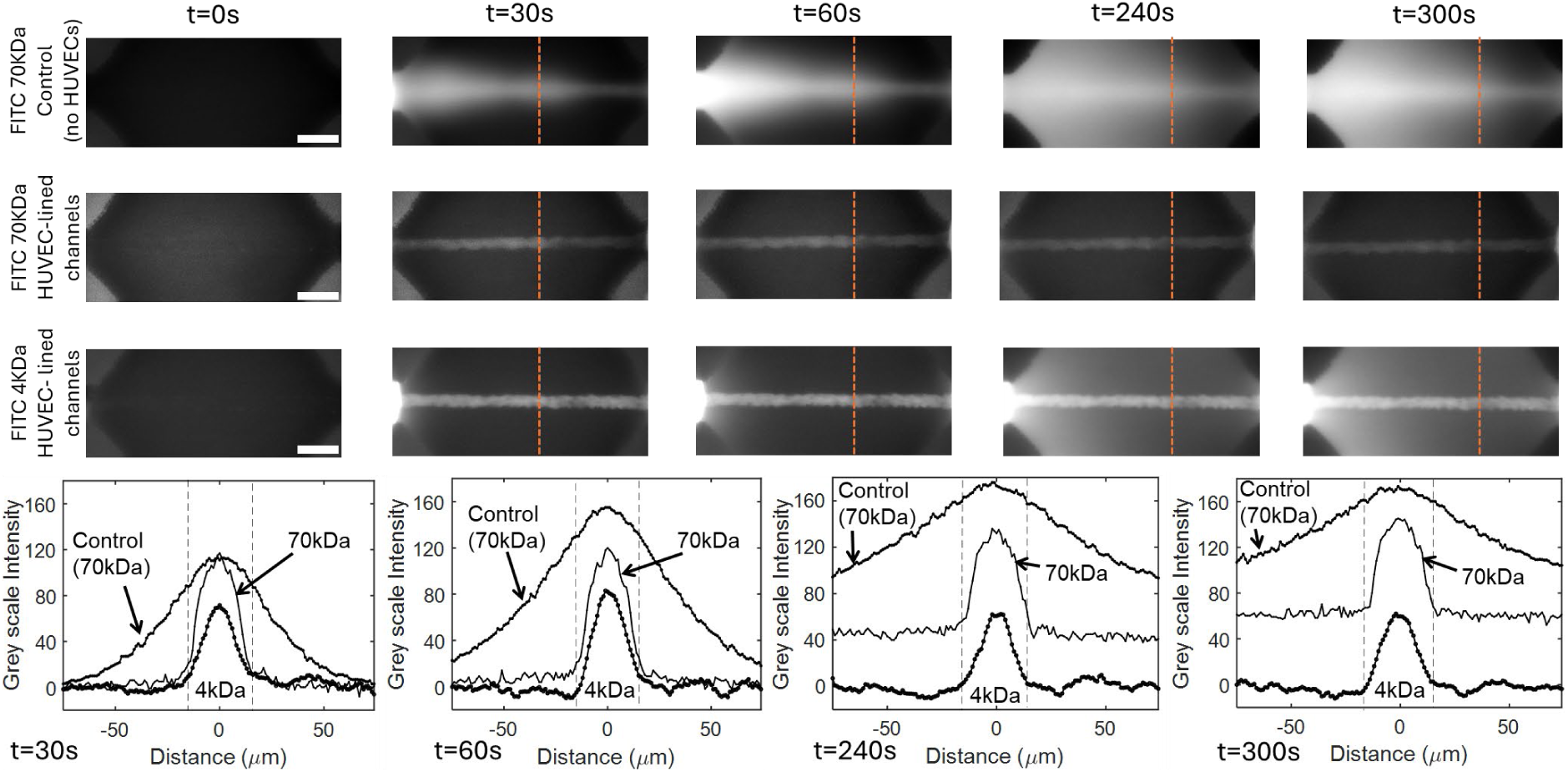
Characterization of patency and barrier function of artificial capillaries using FITC-Dextran (70kDa) for chips with HUVEC-lined microchannels and control samples. In addition, FITC-dextran (4kDa) was also tested for HUVEC-lined microchannels, and changes in intensities across the lumen cross-section was measured. For all experiments, microchannel lumen size = 30µm (n=3). Scale bar: 100 μm

**SI-7.**
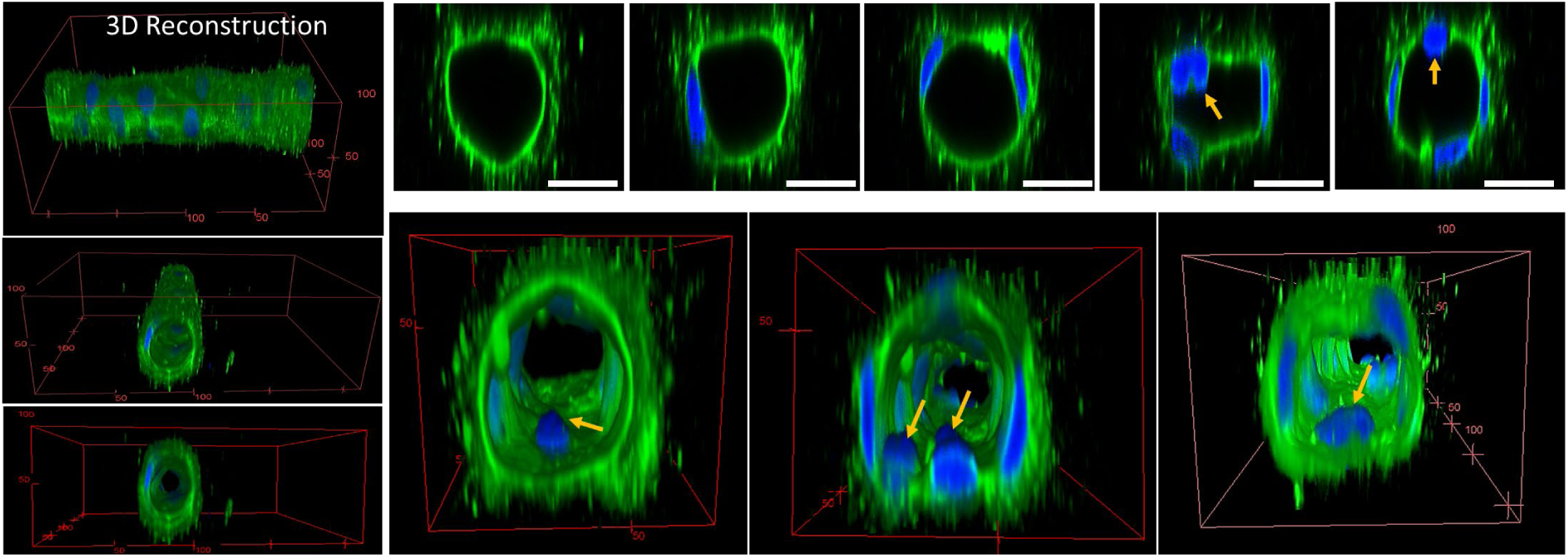
HUVEC-lined lumen showing PECAM staining (CD31, green) and nuclei (blue). Some nuclei can be seen stretching along the sides of the channel while some nuclei protrude into the luminal space (arrow). Scale bar: 25 μm

**SI-8.**
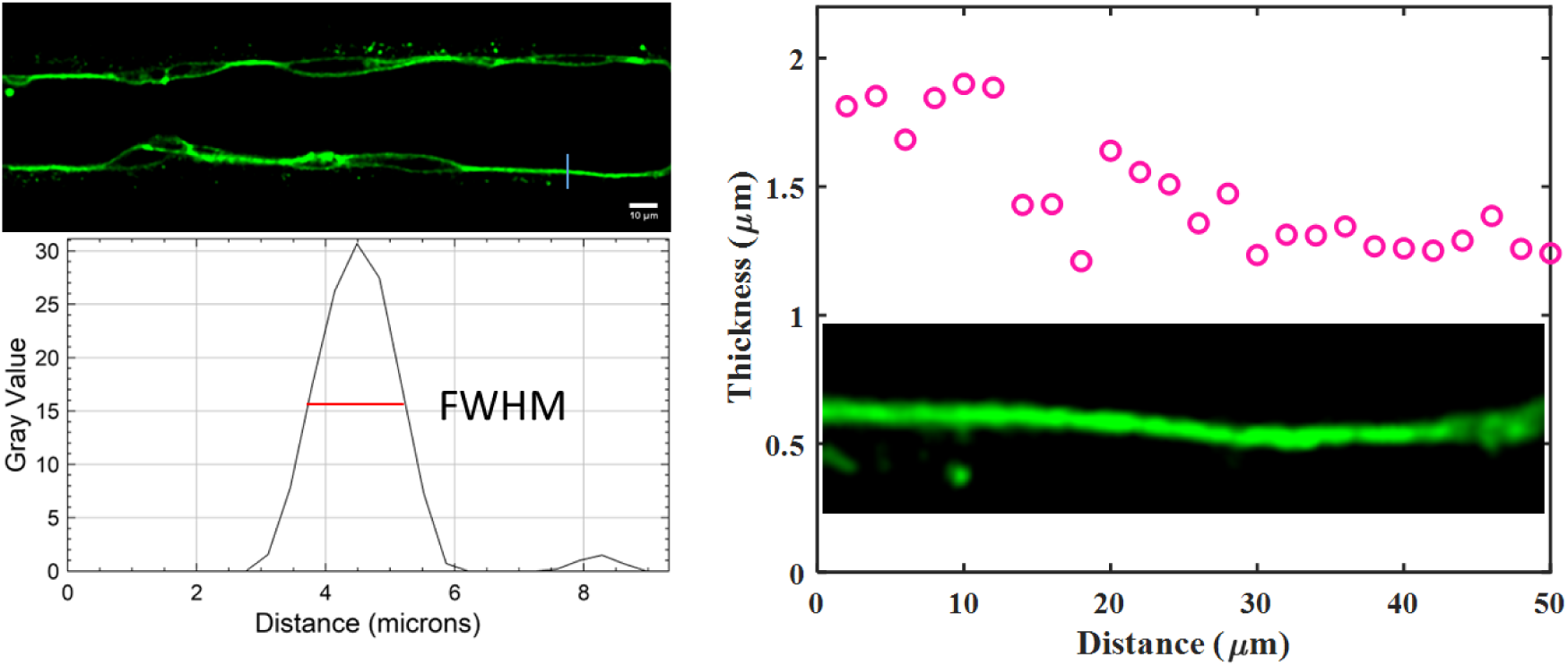
Process to analyze cell thickness in HUVEC-lined microchannels.

**SI-9.**
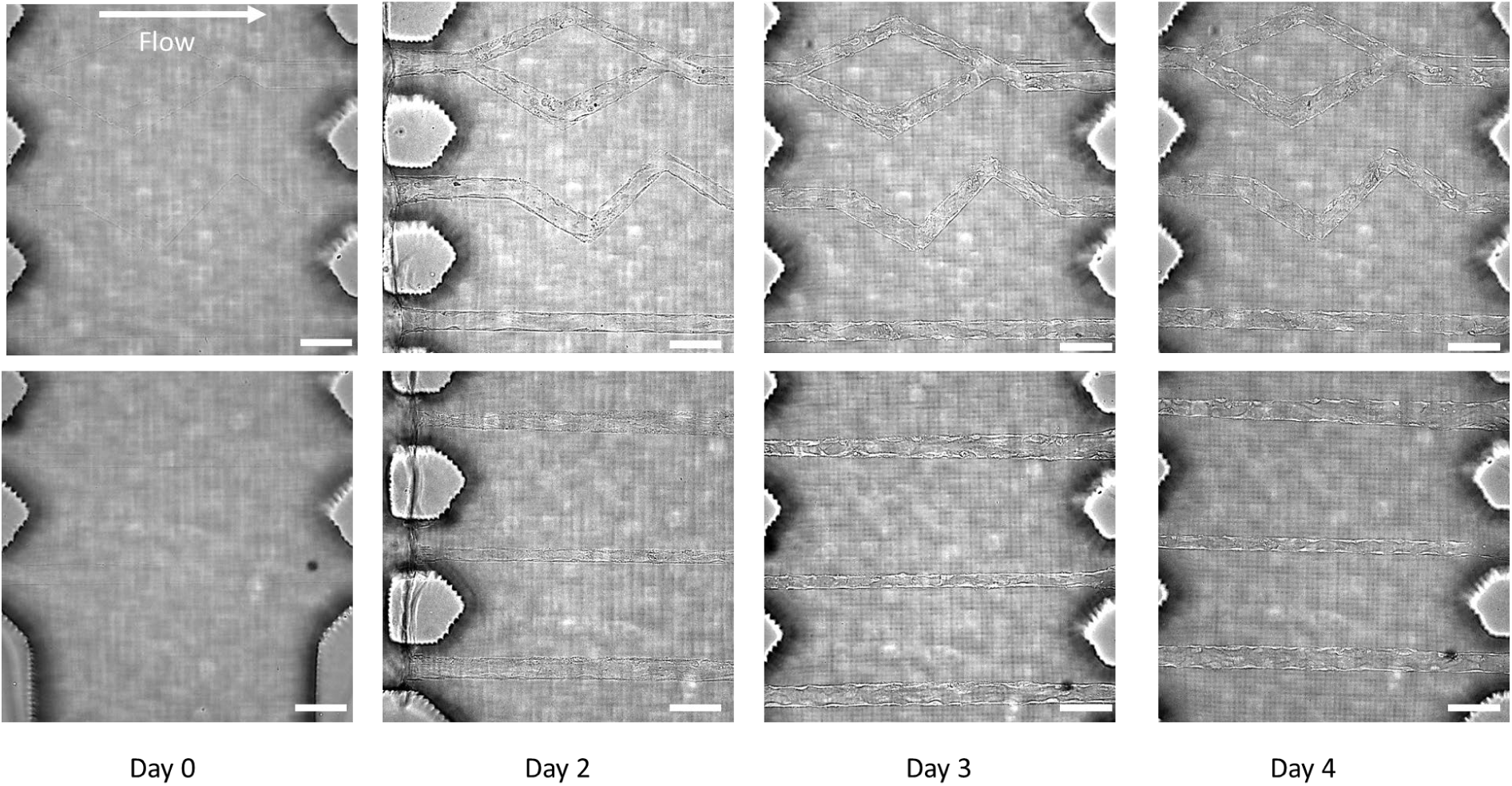
Representative brightfield images with unidirectional flow in microchannels of different geometries (straight, branched, zig-zag) embedded in collagen in Ch#2 of the chip. Scale bar = 100 μm

**SI-10.**
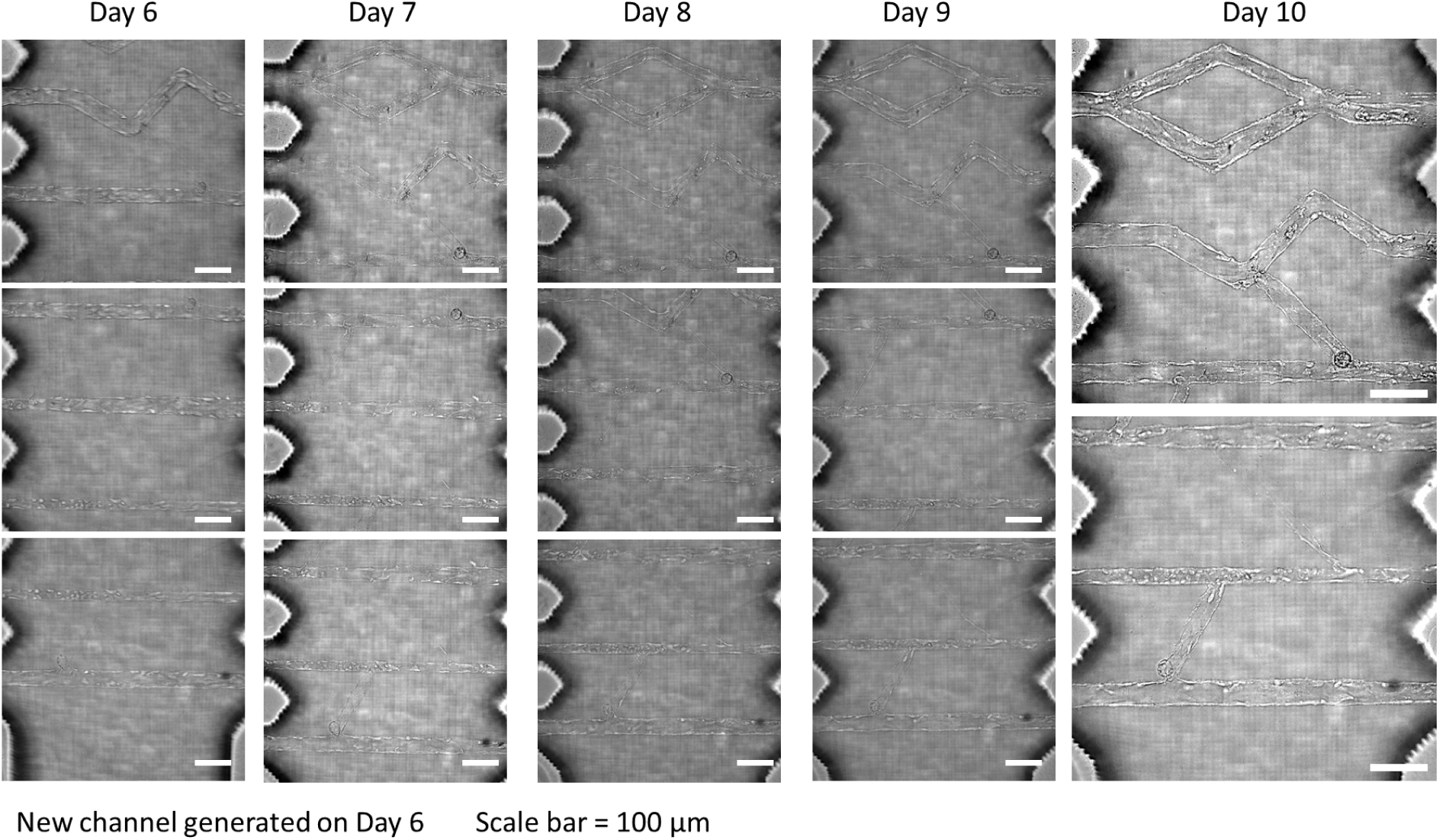
Representative brightfield images with unidirectional flow in channels after new channels were generated by femtosecond laser scanning on Day 6.

**SI-11.**
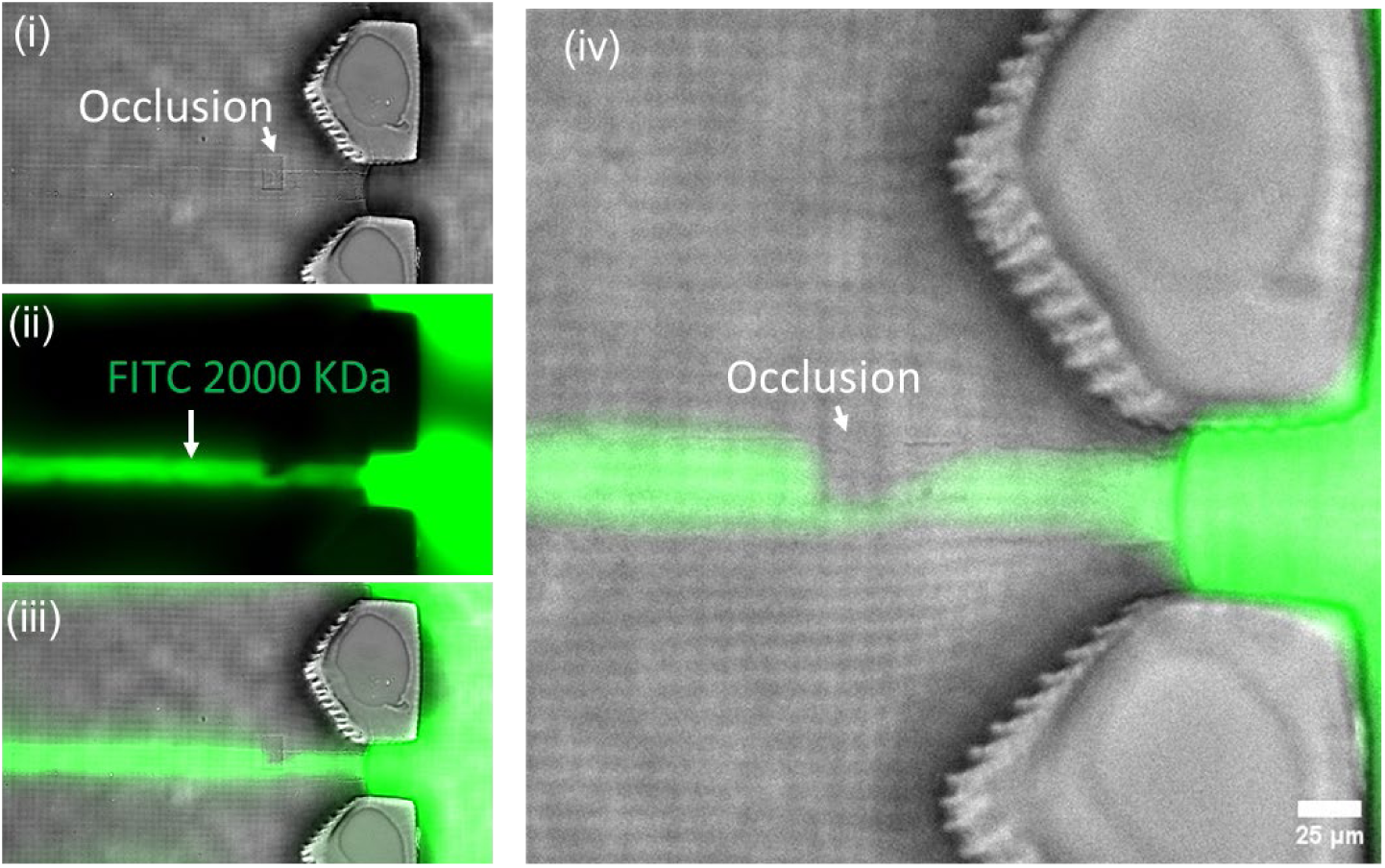
Images showing an addition of a user-defined occlusion in existing endothelialized microchannel via fs-laser crosslinking of GelMA and resulting diversion of flow of perfused FITC-dextran dye solution.

**SI-12:**
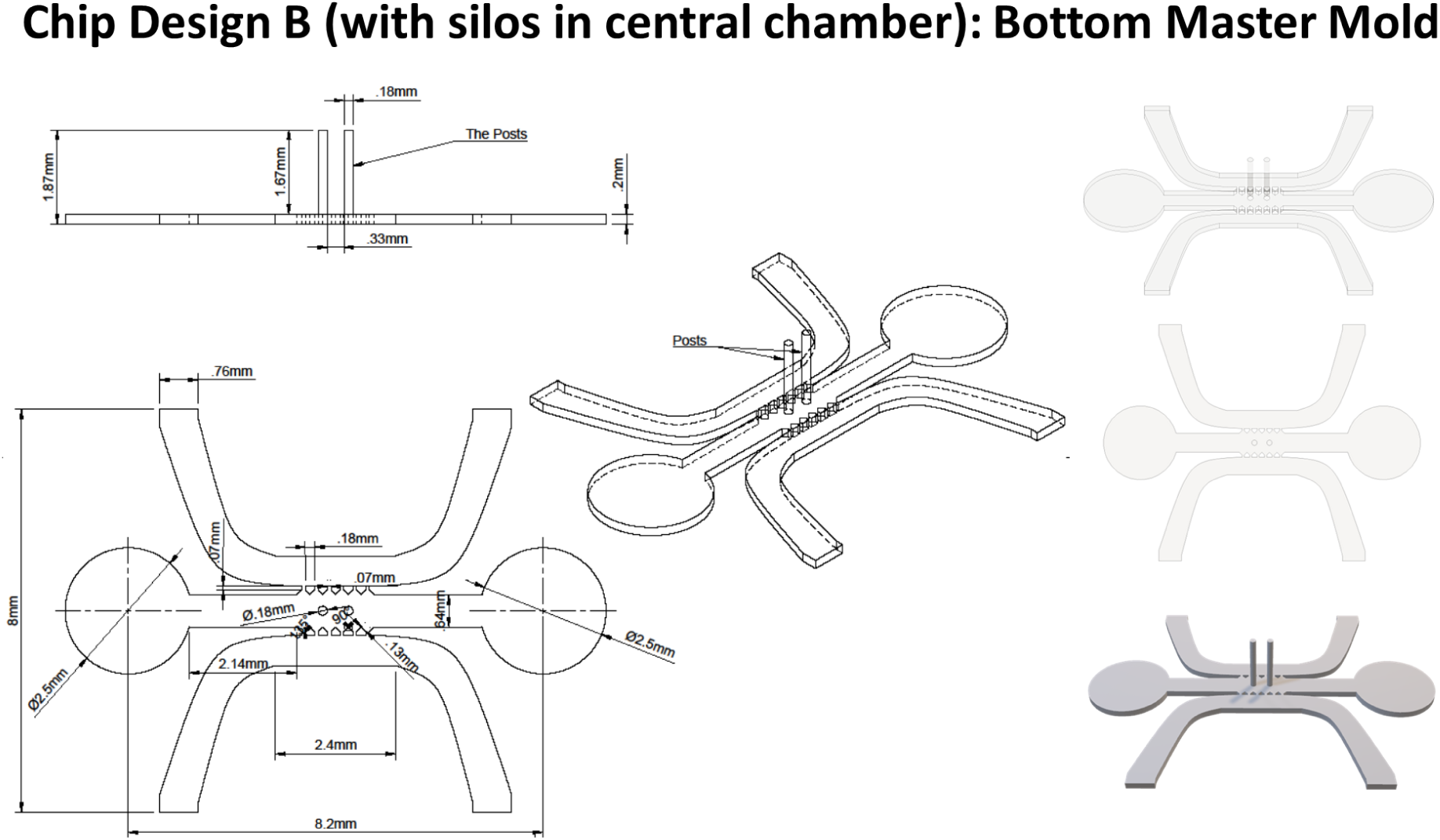
CAD files for the bottom part of a two-part Master Mold for chip design B.

**SI-13.**
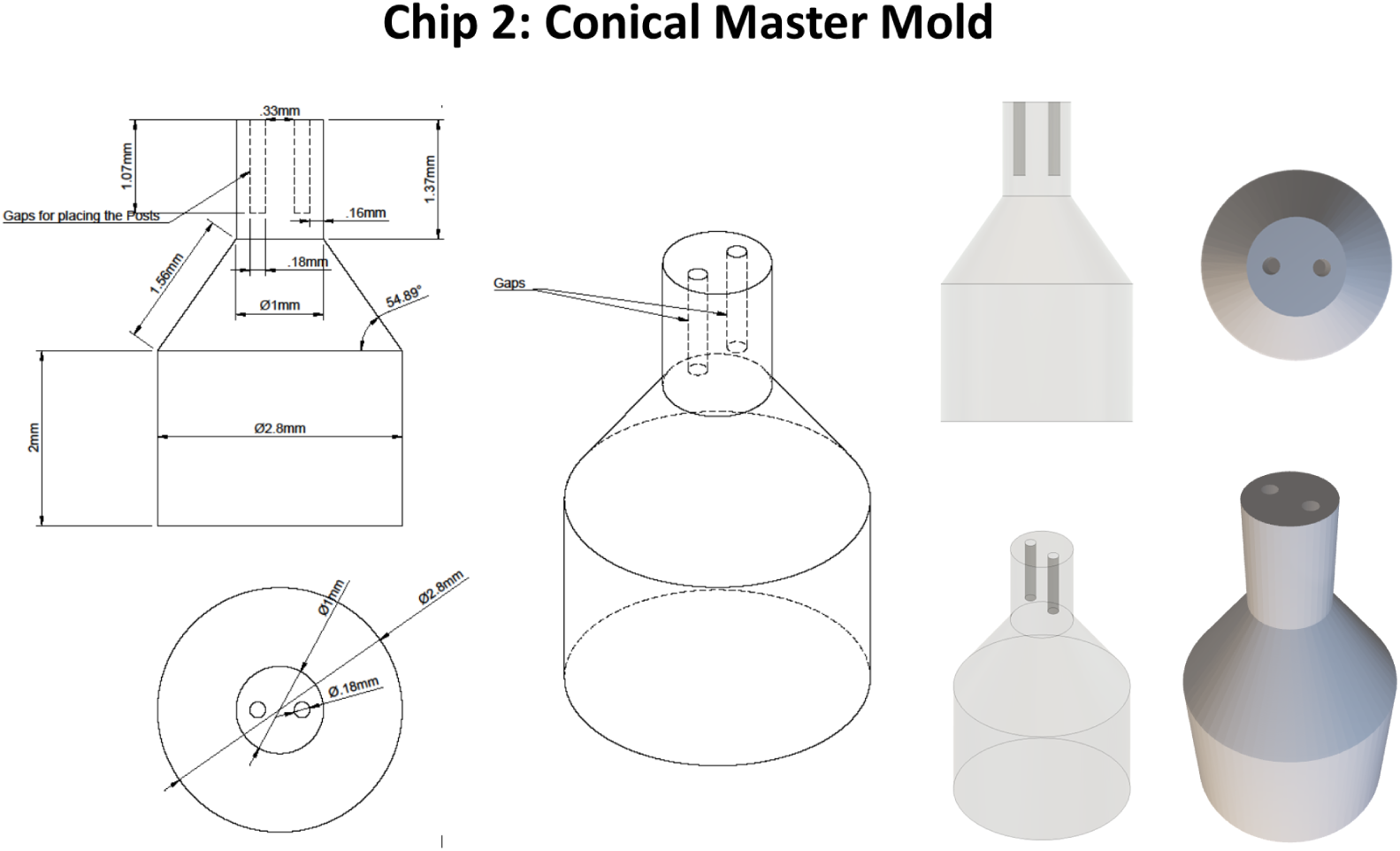
CAD files for the top part of a two-part Master Mold for chip design B

**SI-14.**
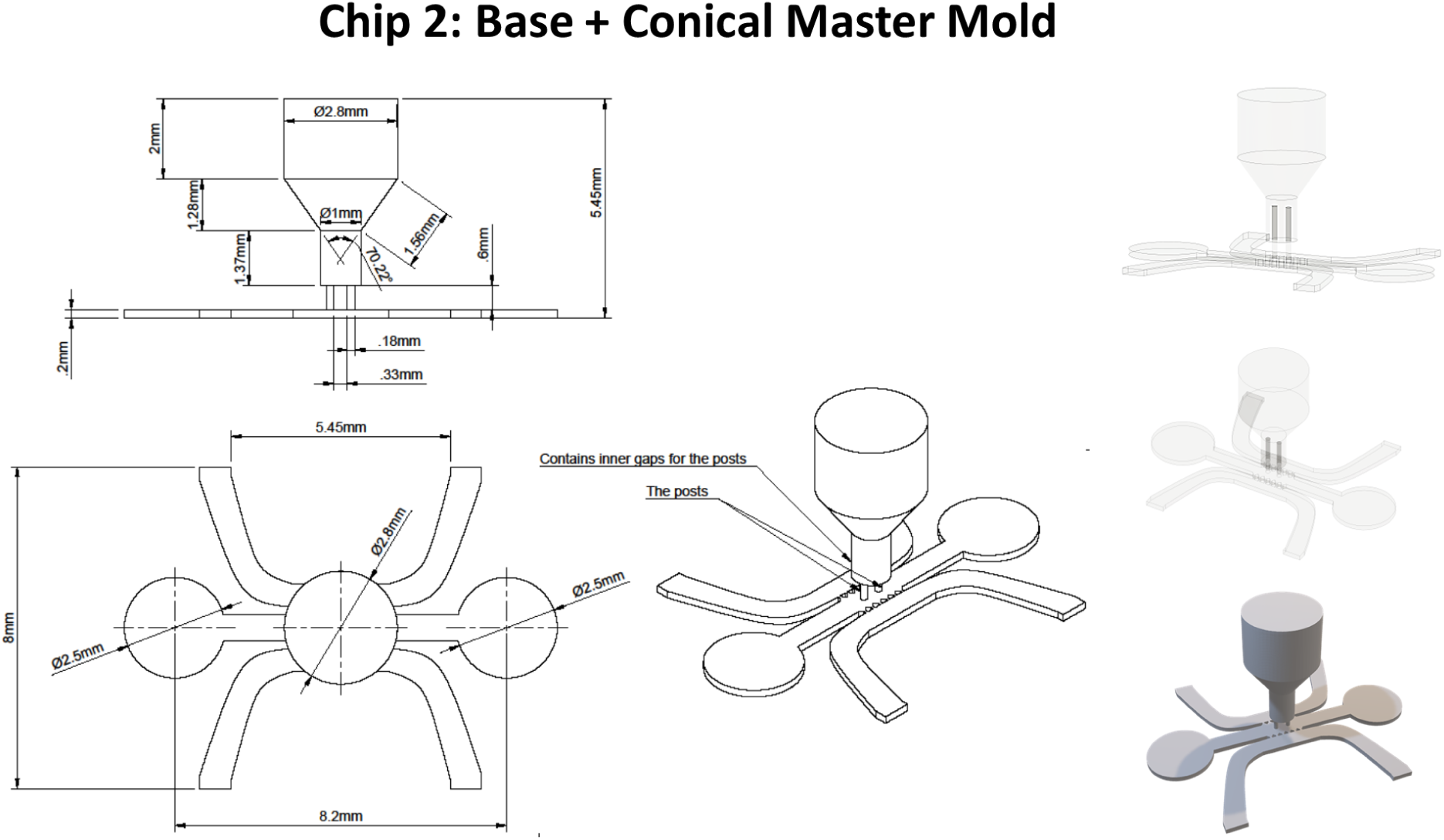
CAD files showing the assembled master mold for chip design B

**SI-15.**
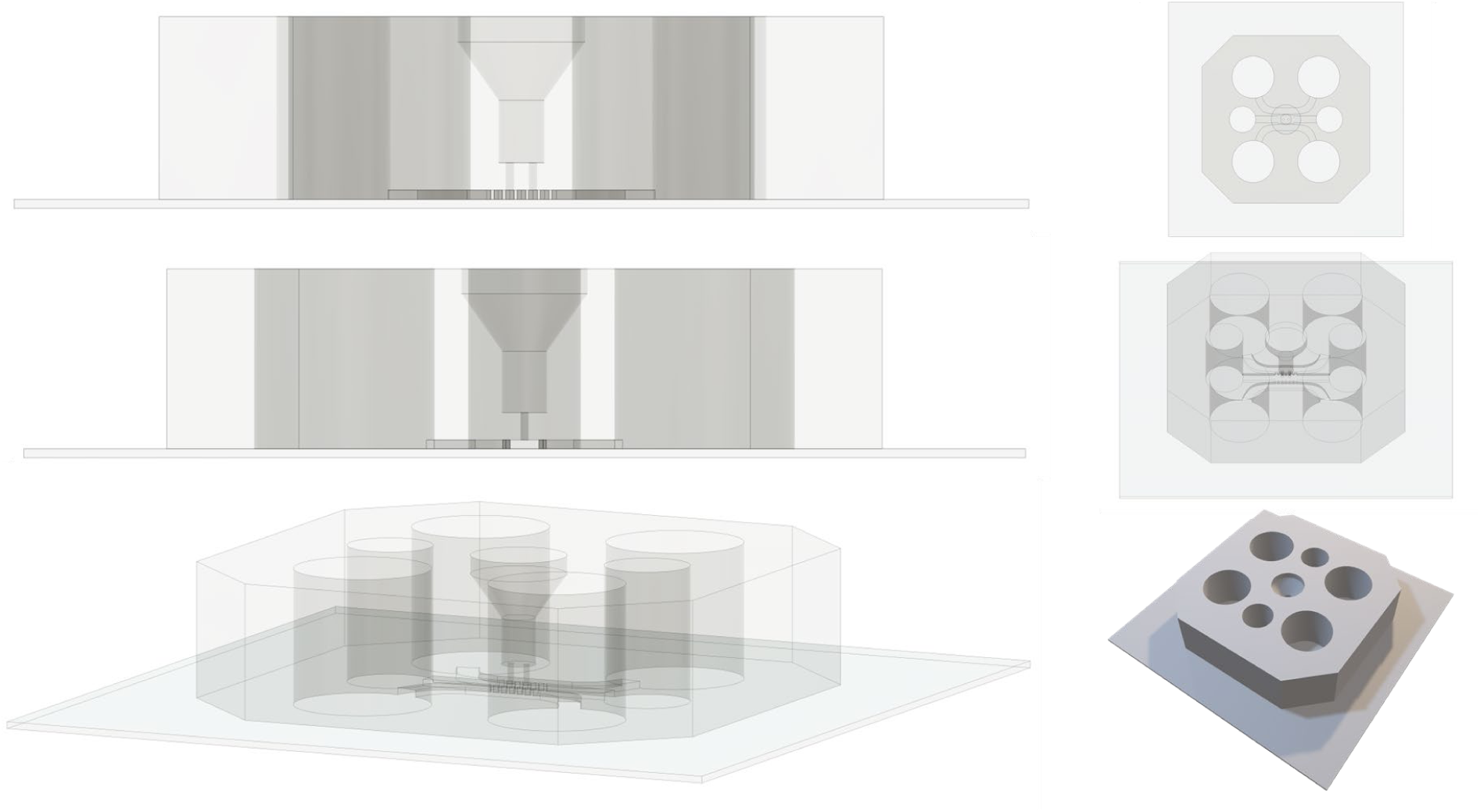
CAD files showing the final PDMS chip for chip design B

**SI-16.**
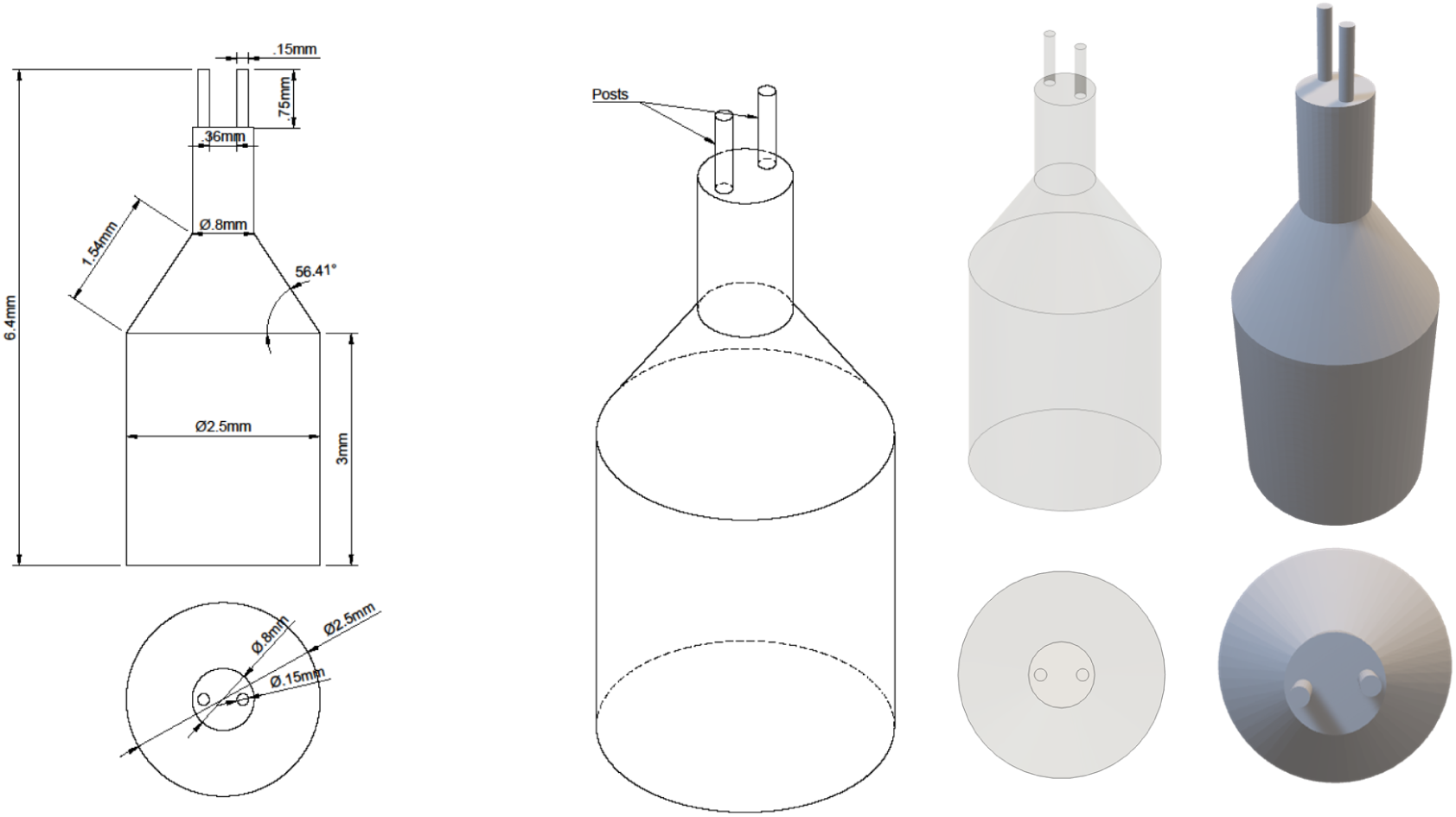
CAD of plug with two pins for chip design B

**SI-17.**
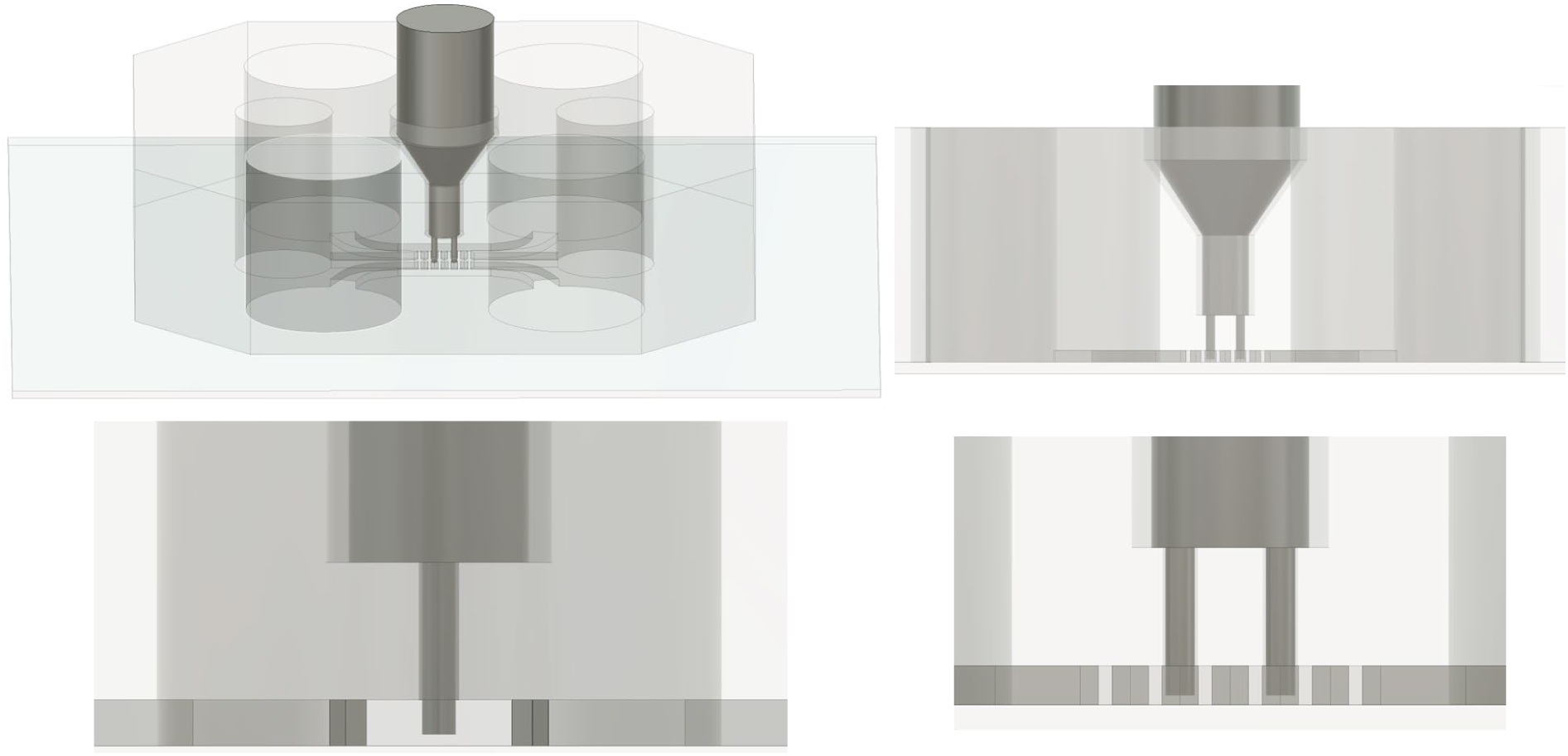
CAD files showing the assembly of plug with two pins into PDMS chip for design B

**SI-18.**
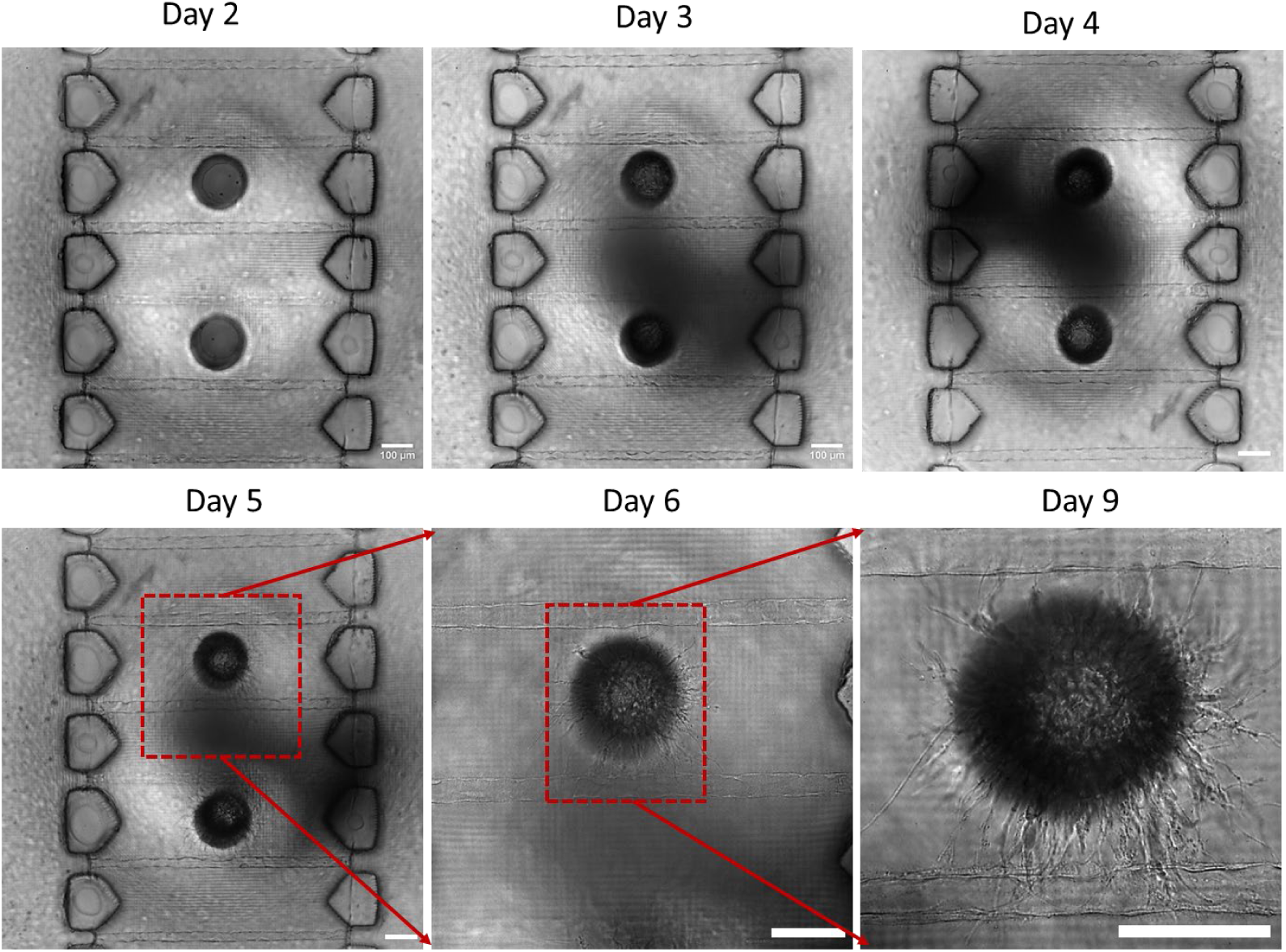
Representative brightfield images showing addition of fibroblasts in silo on Day 3, followed by their migration into collagen matrix, and wrapping around artificial capillaries. Scale bar = 100 μm

**SI-19.**
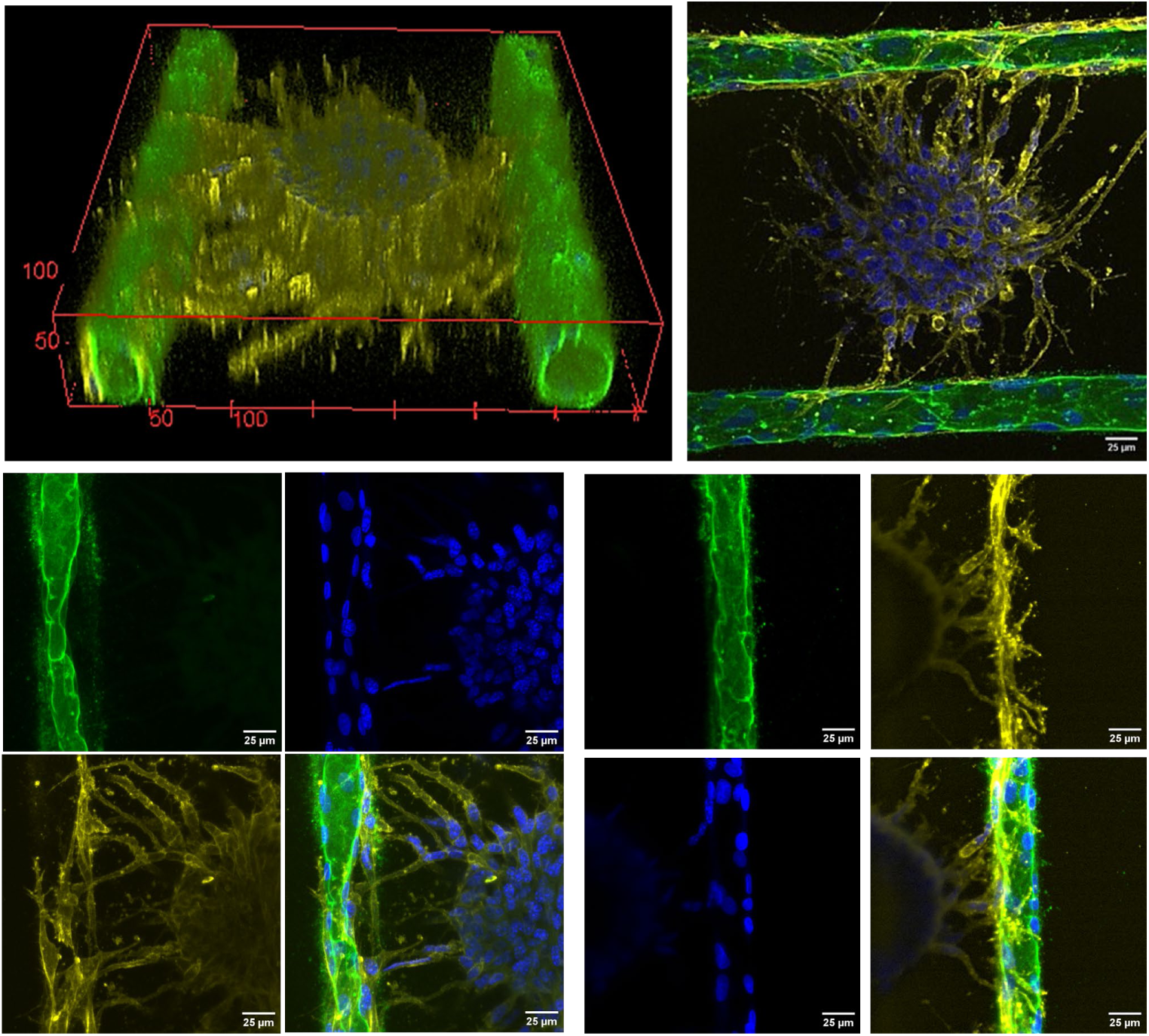
Monitoring Stromal Cells interacting with ECs. 3D reconstruction and fluorescence images showing stromal cells (CD 44: yellow) embedded in collagen extending dendritic projections toward endothelial cell-lined microchannels (CD31:green). Nuclei are stained with DAPI (blue). Stromal cells display clear interactions with the endothelial lining, suggesting physical contact and potential signaling.

**SI-20.**
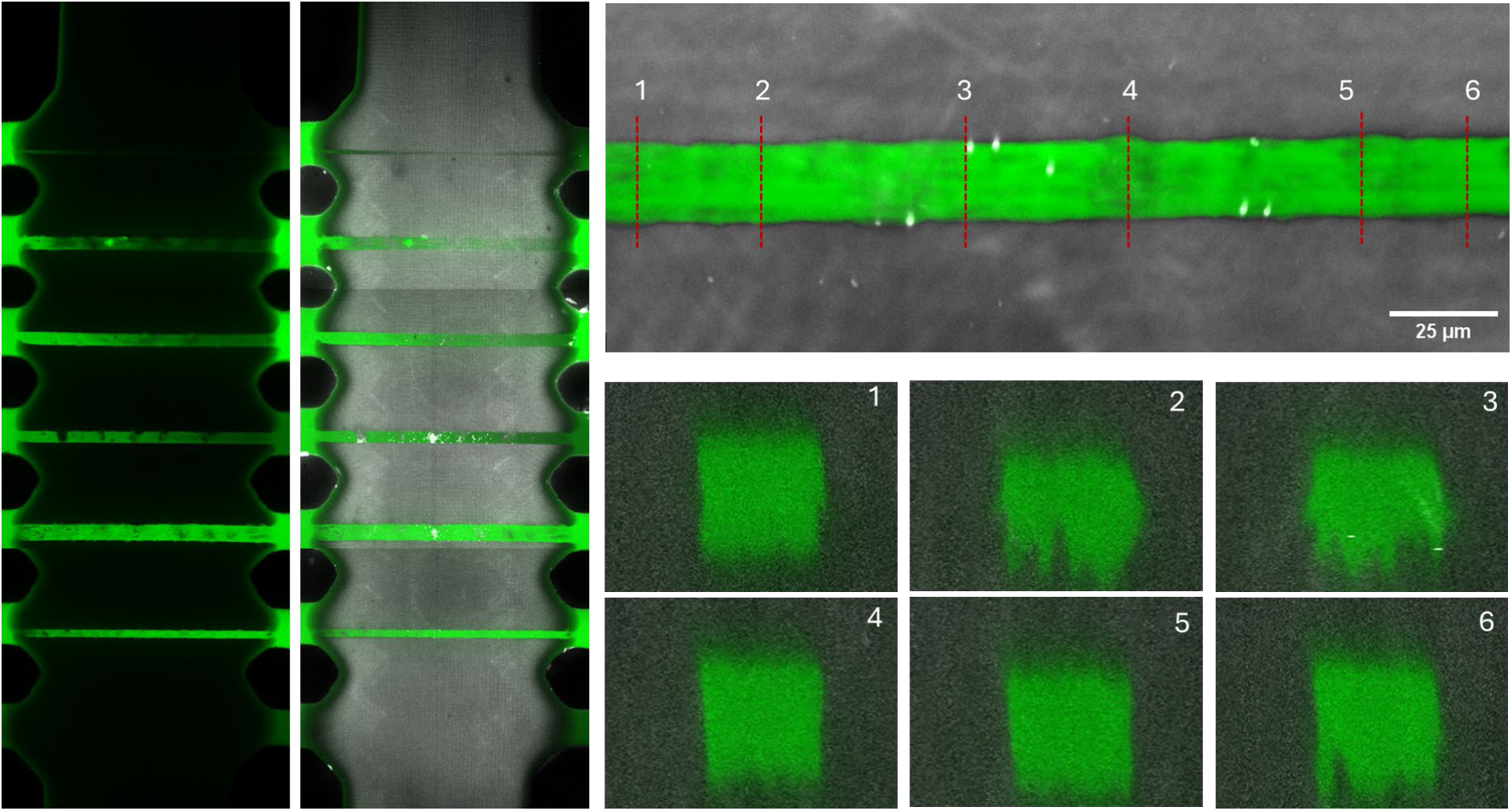
Microchannels in HAMA inside the microfluidic chips

**SI-21.**
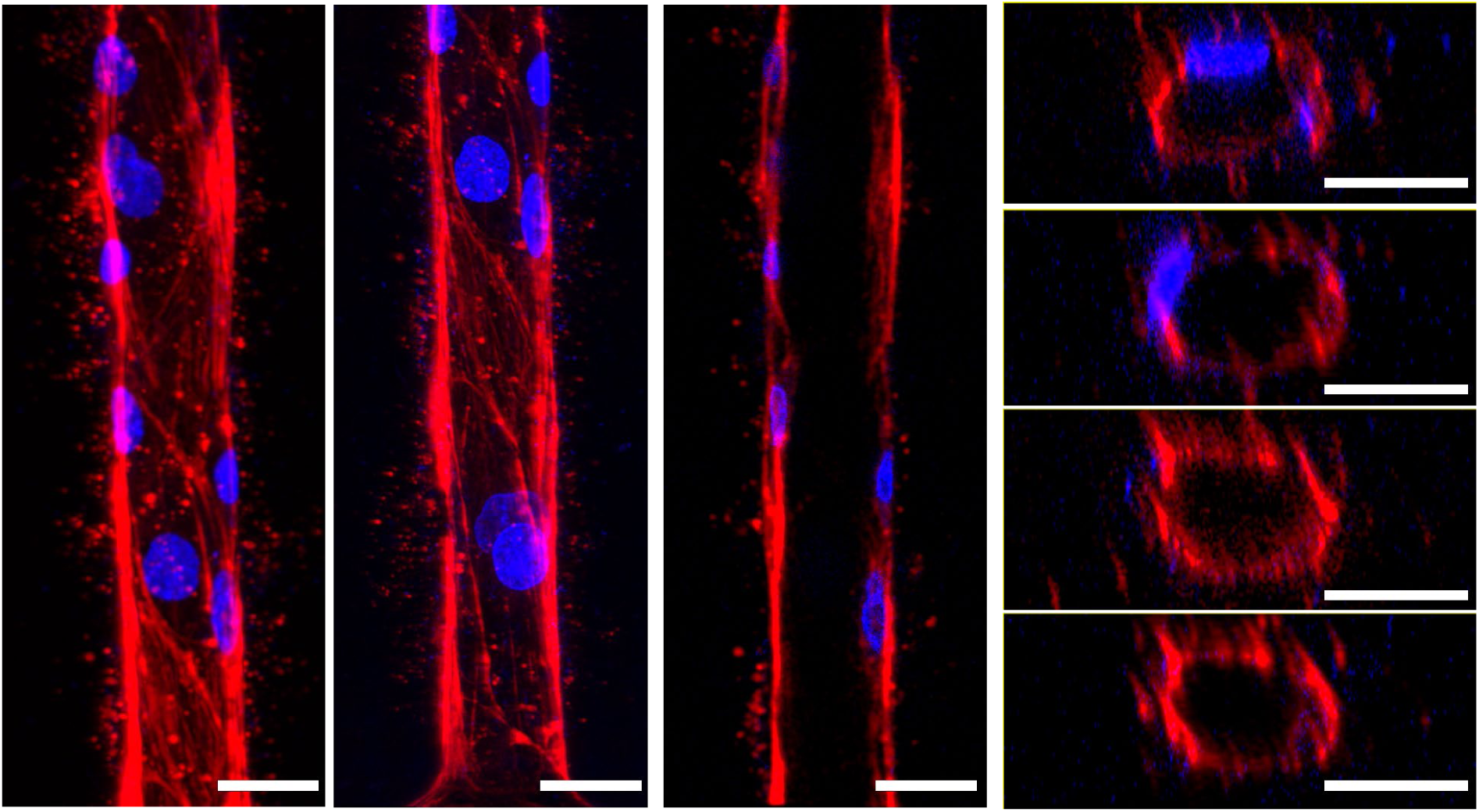
Microchannels lined with Colony Forming Endothelial cells. Scale bar: 25 μm

## Video File Captions

V-1. FITC-dextran (4kDa, green) solution perfused through HUVEC-lined microchannels embedded within collagen matrix.

V-2. FITC-dextran (70kDa, green) solution/dye perfused through acellular microchannels showing diffusion of dye into the collagen matrix.

V-3. Fluorescence z-stack showing adjacent pair of HUVEC-lined microchannels embedded within collagen matrix (f-actin = red; nuclei = blue)

V-4. Fluorescence images showing cross-sections of adjacent pair of HUVEC-lined microchannels embedded with collagen matrix. (f-actin = red; nuclei = blue)

V-5. Fluorescence z-stack showing high-resolution of a single HUVEC-lined microchannel embedded within collagen matrix (CD31 = green; nuclei = blue) Images were acquired using a 40× water-immersion objective.

V-6. Fluorescence images showing cross-section of a single HUVEC-lined microchannel embedded within collagen matrix (CD31 = green; nuclei = blue) Images were acquired using a 40× water-immersion objective.

V-7. Animation of a 3D reconstructed HUVEC-lined microchannel. (CD31 = green; nuclei = blue)

V-8. Animation showing the assembly of top and bottom master molds for chip design 2. Created using Fusion 360 CAD software.

V-9. Animation showing the assembly of the plug with two pins into the 3-chambered PDMS microfluidic chip. Created using Fusion 360 CAD software.

V-10. Fluorescence stack showing cross-sectional views of two HUVEC-lined microchannels located on either side of the central silo with 10T1/2-fibroblasts (stromal cells). (HUVECs are stained using CD31, green; fibroblasts are stained using CD44, yellow; nuclei = blue).

V-11. Fluorescence z-stack showing lateral (XY) views of two HUVEC-lined microchannels located on either side of the central silo with 10T1/2-fibroblasts (stromal cells). (HUVECs are stained using CD31, green; fibroblasts are stained using CD44, yellow; nuclei = blue).

V-12 and V-13. Fluorescence z-stack showing lateral (XY) views of one HUVEC-lined microchannels (lumen size of 8-10 μm) located on either side of the central silo with 10T1/2-fibroblasts (stromal cells). (HUVECs are stained using CD31, green; fibroblasts are stained using CD44, yellow; nuclei = blue)

V-14 and V-15. Fluorescence images showing the cross-sectional views of one HUVEC-lined microchannels (lumen size of 8-10 μm) located on either side of the central silo with 10T1/2-fibroblasts (stromal cells). (HUVECs are stained using CD31, green; fibroblasts are stained using CD44, yellow; nuclei = blue)

V-16. Animation of a 3D reconstructed HUVEC-lined microchannel. (lumen ∼ 8-10 μm) CD31 (green), DAPI (nuclei, blue).

## Notes

### Competing Interest Statement

The authors have declared no competing interest.

